# Social inheritance can explain the structure of animal social networks

**DOI:** 10.1101/026120

**Authors:** Amiyaal Ilany, Erol Akcay

## Abstract

The social network structure of animal populations has major implications for survival, reproductive success, sexual selection, and pathogen transmission of individuals. But as of yet, no general theory of social network structure exists that can explain the diversity of social networks observed in nature, and serve as a null model for detecting species and population-specific factors. Here we propose a simple and generally applicable model of social network structure. We consider the emergence of network structure as a result of social inheritance, in which newborns are likely to bond with maternal contacts, and via forming bonds randomly. We compare model output to data from several species, showing that it can generate networks with properties such as those observed in real social systems. Our model demonstrates that important observed properties of social networks, including heritability of network position or assortative associations, can be understood as consequences of social inheritance.

---

The transition to sociality is one of the major shifts in evolution, and social structure is an important and ever-present selective factor, affecting both reproductive success^1^ and survival^2,3^. Sociality affects individual health, ecological dynamics, and evolutionary fitness via multiple mechanisms in humans and other animals, such as pathogen transmission e.g.^4,5^ and promoting or hindering of particular social behaviors^6,7,8^. The social structure of a population summarizes the social bonds of its members^9^. Hence, understanding the processes generating variation in social structure across populations and species is crucial to uncovering the impacts of sociality.

Recent years have seen a surge in the study of the causes and consequences of social structure in human and animal societies, based on theoretical and computational advances in social network analysis ^10,11,12,13,14,15,16^. The new interdisciplinary network science provides many tools to construct, visualize, quantify and compare social structures, facilitating advanced understanding of social phenomena. Researchers studying a variety of species, from insects to humans, have used these tools to gain insights into the factors determining social structure ^17,18,19,13^. Using social network analysis provided evidence for the effects of social structure on a range of phenomena, such as sexual selection^20^ and cultural transmission^21,22^.

At the same time, most applications of social network analysis to non-human animals have been at a descriptive level, using various computational methods to quantify features of social structure and individuals’ position in it. These methods, combined with increasingly detailed data “reality mining”^23^ about social interactions in nature, provided valuable insights about the complex effects of social interaction on individual behaviors and fitness outcomes. Yet, we still lack a comprehensive theory that can explain the generation and diversity of social structures observed within and between species. There have been only a few efforts to model animal social network structure. Notably, Seyfarth^24^ used a generative model of grooming networks based on individual preferences for giving and receiving grooming, and showed that a few simple rules can account for complex social structure. This model and related approaches e.g.,^25^ have been very influential in the study of social structure and continue to drive empirical research. At the same time, they mostly focused on primates and were geared towards specific questions such as the effects of relatedness, social ranks, or ecological factors in determining social structure.

Independently, a large body of theoretical work in network science aims to explain the general properties of human social networks through simple models of how networks form. Yet these models tend to focus either on networks with a fixed set of agents^26^, or on boundlessly growing networks^27^, with few exceptions^28,29^. These network formation models therefore have limited applicability to animal (and many human) social groups where individuals both join (through birth of immigration) and leave (through death or emigration) the network. Furthermore, most work in network science concentrates on the distribution of number of connections individuals have (the degree distribution). Models that fit the degree distribution of real-world networks tend to be a poor fit to other important properties, notably the tendency of social connections to be clustered^30,27^, i.e., two individuals to be connected with each other if they are both connected to a third individual. Real-world human and animal networks exhibit significantly more clustering than random or preferential attachment models predict^27,13^.

Simple generative models of complex systems have been highly useful in other fields, such as metabolic networks^31^ and food webs^32^, but there has been little effort to build such models applicable to animal social networks. In this paper, we provide a widely applicable network formation model based on simple demographic and social processes. Our model assumes a simple neutral demography and focuses on a central social process that is in operation in many social species: the “inheritance" of social connections from parents. This central component of our model is based on the observation that in many species with stable social groups, individuals interact with the social circle of their parents. This is essentially the case in all mammals, where newborns stay close to their mothers until weaning, but also found in many other taxa, such as birds^33^, fish^34^, and arthropods^35^. After positively interacting with the parents’ social contacts, young individuals are likely to form social bonds with these conspecifics, as was found in African elephants, *Loxodonta africana* ^36^.

We demonstrate that this simple social inheritance process can result in networks that match both the degree and local clustering distributions of real-world animal social networks, as well as their modularity (which measures the strength of division of a network into modules, or subgroups). We also show that social heritability of connections can result in the appearance of genetic heritability of individual social network traits, as well as assortativity in the absence of explicit preference for homophily. Our approach highlights commonalities among groups, populations, and species, and uncovers a general process that underlies variation in social structure.

## Results

Our departure point is the model by Jackson and Rogers^27^, in which “role models” in a network introduce their new contact to their other contacts. This model can reproduce many attributes of large-scale human social networks. Similar models reconstruct the structure of other systems, such as protein interaction networks^37^, and the World Wide Web^38^. However, Jackson and Rogers’ model (like most other models in this family) is based on a constantly growing network with no death or emigration of agents and their results hold asymptotically for very large networks. Since we are interested in small-scale animal networks that do not grow unboundedly, we model a population where existing individuals die and get replaced at an equal rate with newborn individuals^28^ (see SI 8 for results for slowly growing and shrinking networks). We model binary and undirected networks, so implicitly assume social bonds are neutral or cooperative, but our model can be extended to weighted networks that describe the strength of each social bond, and directed ones, such as agonistic networks.

Consider a social group of size *N*. Suppose that each time step, an individual is born to a random mother, and one individual is selected to die at random. With probability *p_b_*, the newborn will meet and connect to its mother (generally, *p_b_* will be close to one, but can be low or zero in species such as some insects, where individuals might not meet their mothers). A crucial component of our model is the general assumption that the likelihood of a newborn A connecting with another individual B depends on the relationship between A’s mother and B: the probability A will connect to B is given by *p_n_* if A’s mother is connected to B, and *p_r_* if not (*n* and *r* stand for neighbor and random node, respectively; Figure **1**). Hence, *p_n_* is the probability an offspring “inherits” a given connection of its parent. Here, we focus on the case A always connects to its mother (*p_b_* = 1), but the model can be extended to include a lower probability to connect to the contacts of A’s mother if A does not connect to its mother, when *p_b_* < 1. If *p_n_* > *p_r_*, the population exhibits a tendency for clustering, a well-established and general phenomenon in social networks^39,13^. In the Supporting Information section we present an extension of this basic model to account for two sexes, where only females reproduce. We show that if newborns are likely to copy only their mothers, the resulting social network is similar.

**Figure 1:**
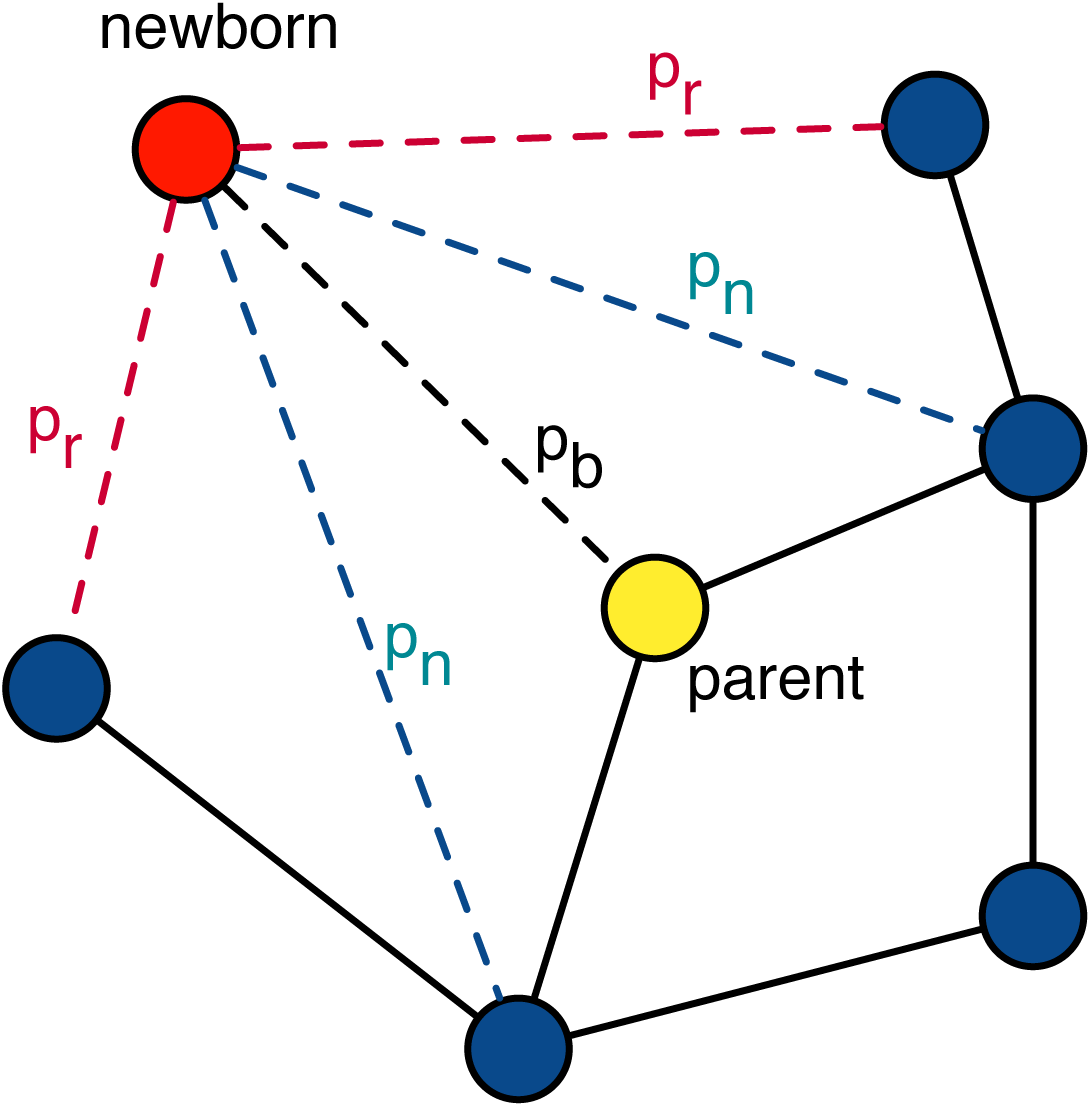
Graphical illustration of the model. A newborn individual is connected to its parent with probability *p_b_*, to its parent’s connections with probability *p_n_*, and to individuals not directly connected to its parent with probability *p_r_*.

We simulated social network dynamics to test how social inheritance and stochastic social bonding affect network structure, heritability, and assortativity (see Methods for simulation details). We also provide analytical expressions for the degree distribution, and approximations for mean degree and mean local clustering coefficient in the Methods section and in the Supplementary Information (SI; section SI 1). For all of our numerical results, we assume *p_b_* = 1. As expected, the network density (the number of edges out of all possible edges) depends on *p_n_* and *p_r_*. The mean clustering coefficient, a measure of the extent to which nodes tend to cluster together, also depends on these parameters, but not monotonically; high levels of clustering were observed in simulations with low or high *p*_r_, but not at intermediate levels (Fig. **2**). We also tested how changes in network size, caused by increased or decreased probabilities of death during the simulations, affected its properties. These tests did not provide a general conclusion, but suggested that the network structure might be moderately influenced by whether the network is growing or not (see SI 8).

**Figure 2:**
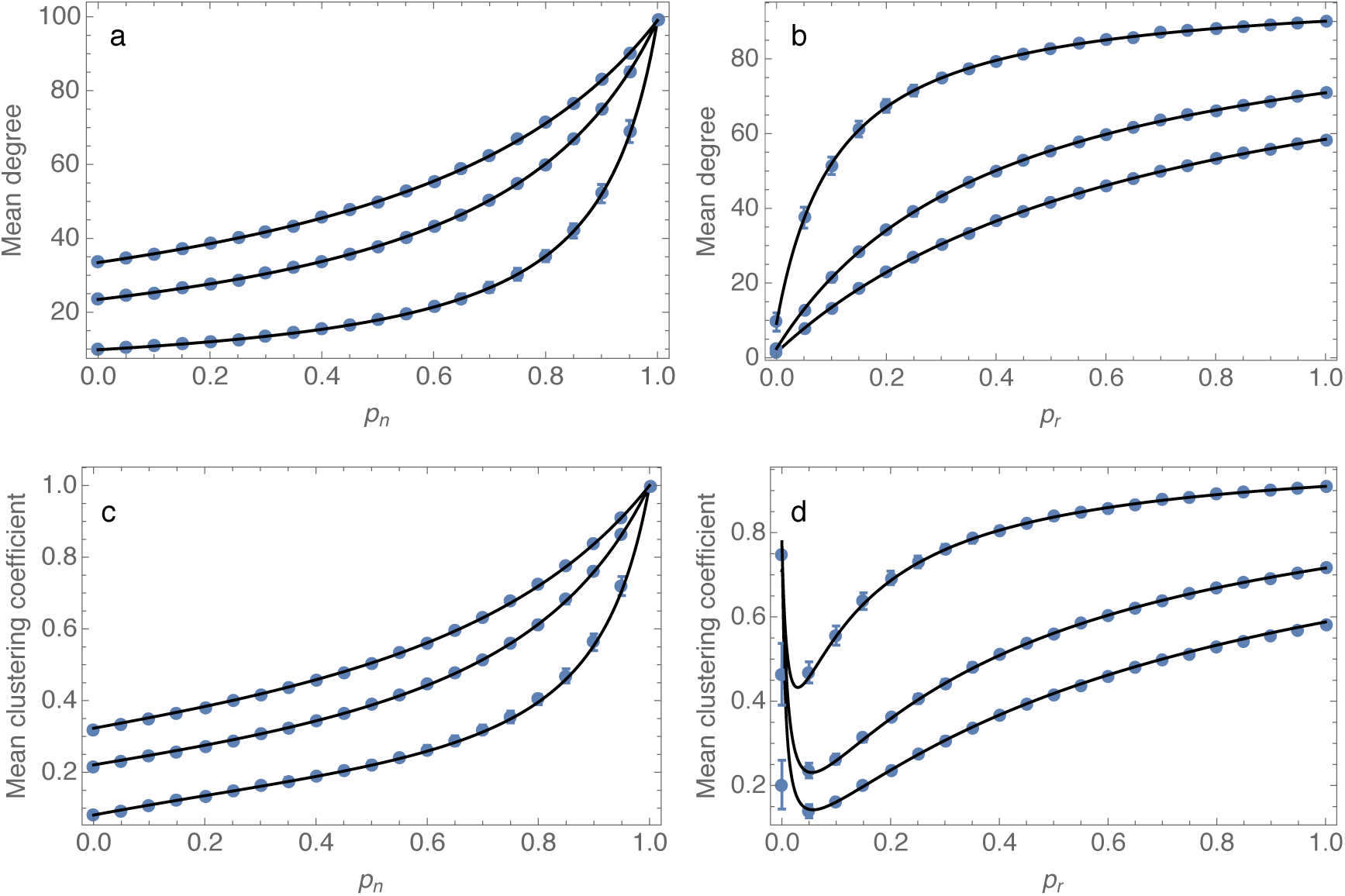
Mean degree and clustering coefficient as a function of model parameters. The dependency of mean degree (top row) and clustering coefficient (bottom row) on social inheritance, *p_n_* (left column), and probability of random bonding, *p_r_* (right column). In each panel, the black curves depict our analytical approximation while the blue dots with error bars are mean and standard deviation of 50 replicate runs. For the two panels on the left, the curves correspond to, from top to bottom, *p_r_* = 0.5, 0.3 and 0.1; for the two panels on the right, from top to bottom, *p_n_* = 0.9, 0.6 and 0.3. For all panels, network size *N* = 100; the simulations were initiated with random networks and run for 2000 time steps.

We then compared the output of our model with observed animal social networks of four different species, namely spotted hyena (*Crocuta crocuta*^13^), rock hyrax *(Procavia capensis*^40^), bottlenose dolphin (*Tursiops* spp.^41^), and sleepy lizard (*Tiliqua rugosa*^42^). We used two independent ways to estimate model parameters using data from each of the four species: a computational dimensionality reduction approach (partial least squares regression, PLS) and analytical approximations for the mean degree and local clustering coefficients (see Methods). When we run our model using *p_n_* and *p_r_* estimated from the data using either method, we recapture the distributions of degree and local clustering coefficient, as well as the network modularity. Figure **3** illustrates that our model of social inheritance can produce networks with realistic social structure (see SI 4 for fitting the two-sex model to observed networks). Our model’s good match of local clustering distributions distinguishes it from other network growth models, based on assortative or generalized social preferences, as well as the preferential attachment models that are popular in network science^27^ (see also SI 6 and SI 7). Furthermore, our model generated networks with realistic modularity values (see SI, figure S5). The values we found suggest that social inheritance is stronger in hyena and hyrax than in dolphins and sleepy lizards (Table 1).

**Figure 3:**
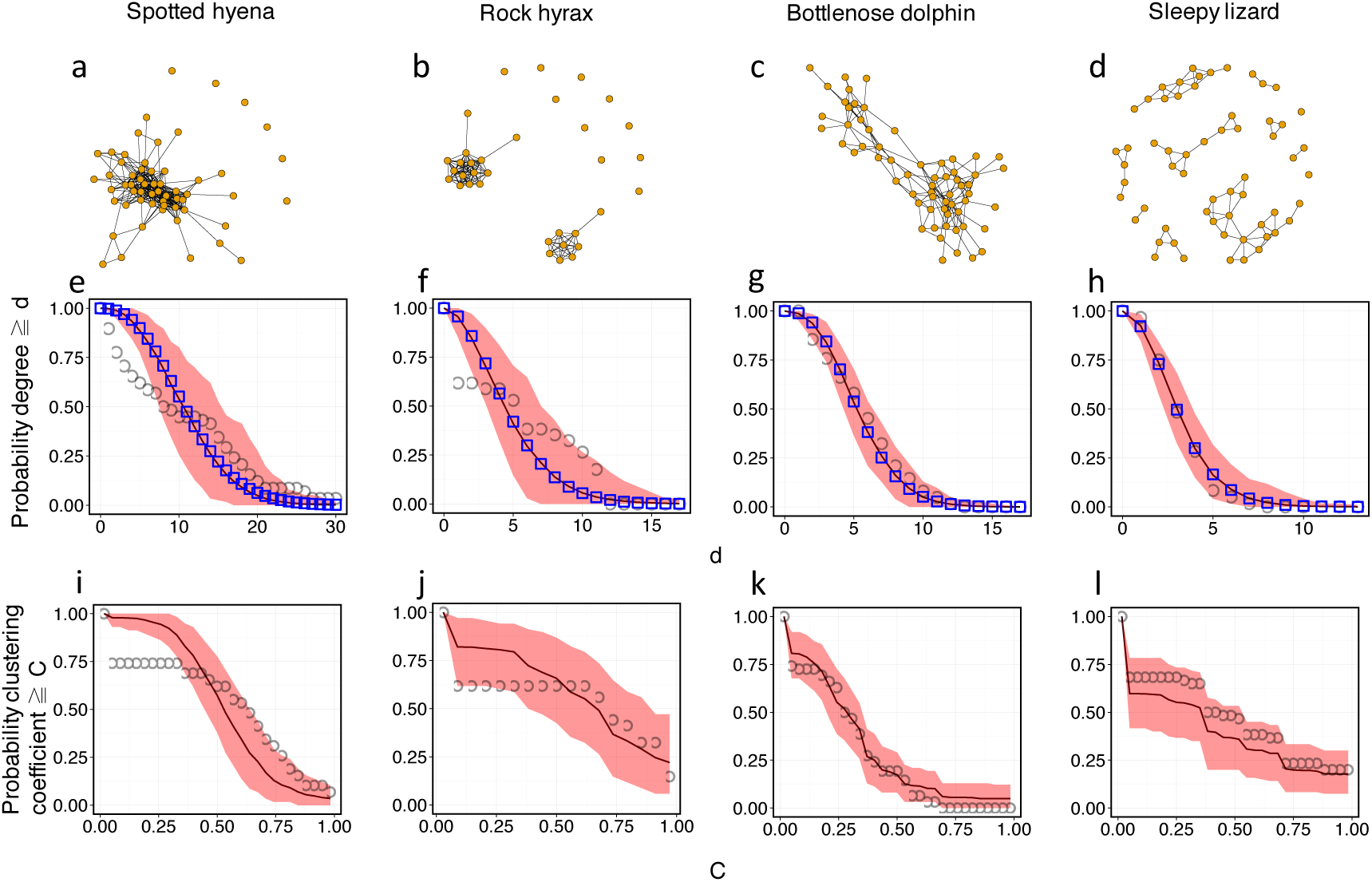
Social inheritance captures essential properties of animal social networks in the wild. This figure shows that our model can account for the degree and clustering coefficient distributions of observed networks in four species. Upper row: observed networks. Middle row: Cumulative degree distributions of observed and simulated networks (d stands for degree). Lower row: Cumulative local clustering coefficient distributions of observed and simulated networks (C stands for clustering coefficient). Black circles represent observed values. Blue squares in the middle row depict mean-field estimation for the degree distribution. The red curve denotes mean distribution for 500 simulated networks (2000 simulation steps) with the species-specific *p_n_* and *p_r_* values estimated using partial least squares regression (values given in Table 1; see Methods for more on the estimation procedure), whereas light red area depicts 95% confidence intervals.

**SI Figure 1:**
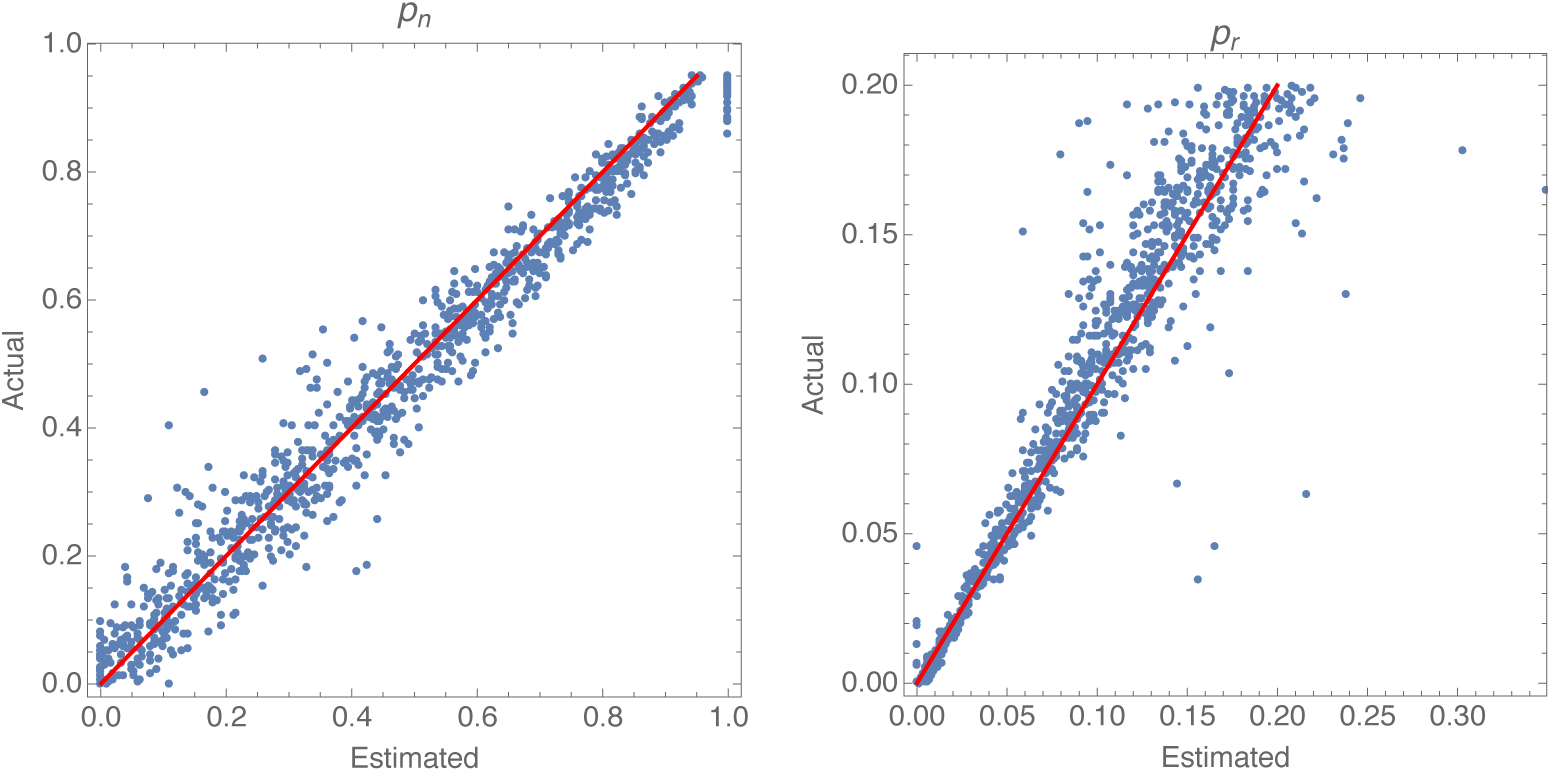
Estimation of model parameters using the analytical approximation. The analytical approximations for the mean degree and clustering coefficient allow us to estimate network parameters with good accuracy. The two panels show for 1000 simulated networks (*N* = 100) the actual *p_n_* (left) and *p_r_* (right) values plotted against the values estimated from our analytical approximation. In each panel, the red line depicts the 1:1 relationship. For each simulation, the *p_n_* and *p_r_* values were drawn randomly from a uniform distribution on [0,0.95] and [0,0.2], respectively. We initialized each simulation with a random network and ran it for 2000 steps. We calculated the mean degree and local clustering coefficients for the resulting network. We used these values to numerically solve equations (1) and (SI-10) for *p_n_* and *p_r_*, to obtain the estimates.

**SI Figure 2:**
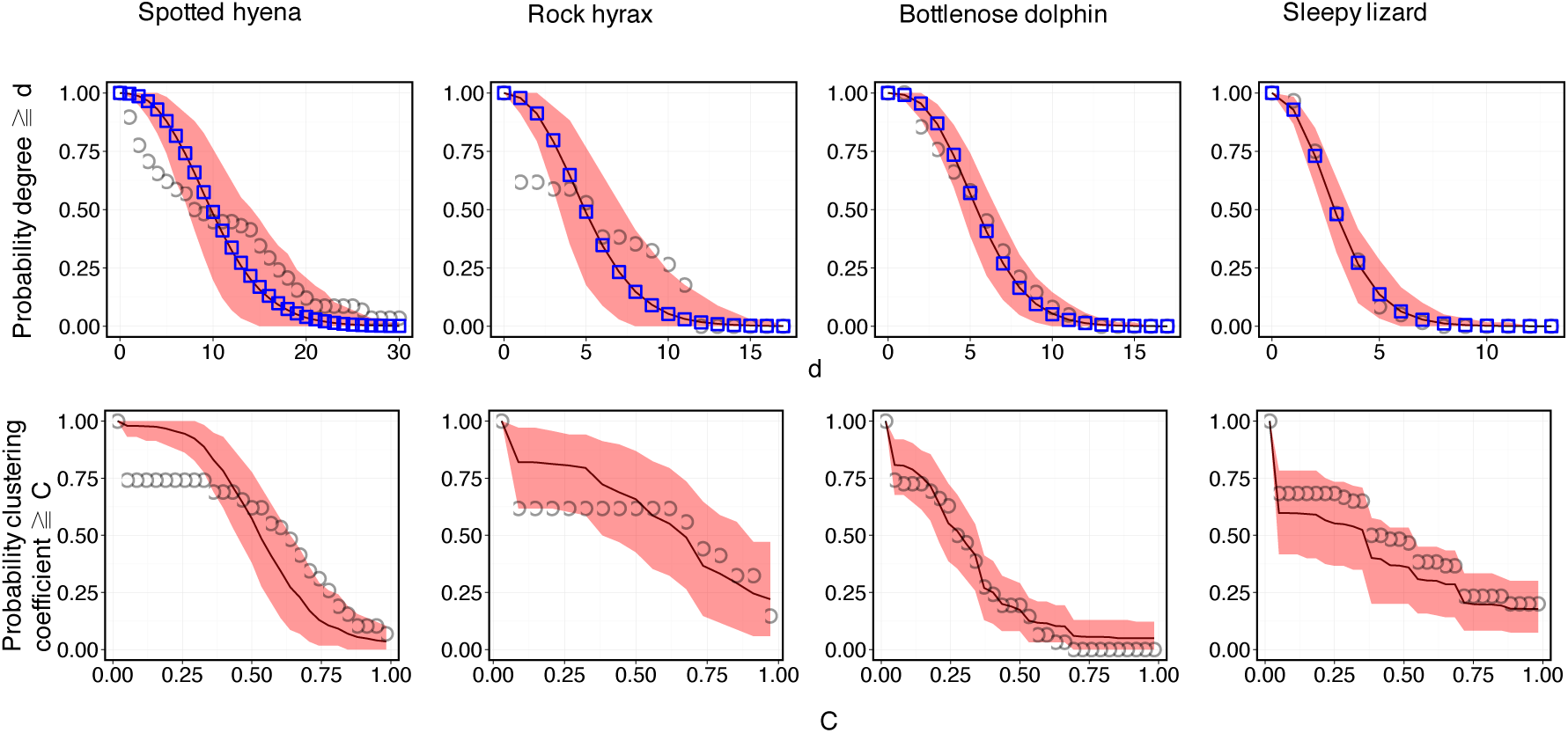
Fitting model to data using the analytical approximation. This figure compares networks generated using parameters estimated from the analytical approximation to the data from four species. Upper row: Cumulative degree distributions of observed and simulated networks. Lower row: Cumulative clustering coefficient distributions of observed and simulated networks. Black circles represent observed values. Blue squares in the upper row depict mean-field estimation for the degree distribution. Red line notes mean values for 500 simulated networks (2000 simulation steps) with the same species-specific *p_n_* and *p_r_* values (given in Table 1), whereas light red area depicts 95% confidence intervals.

**SI Figure 3:**
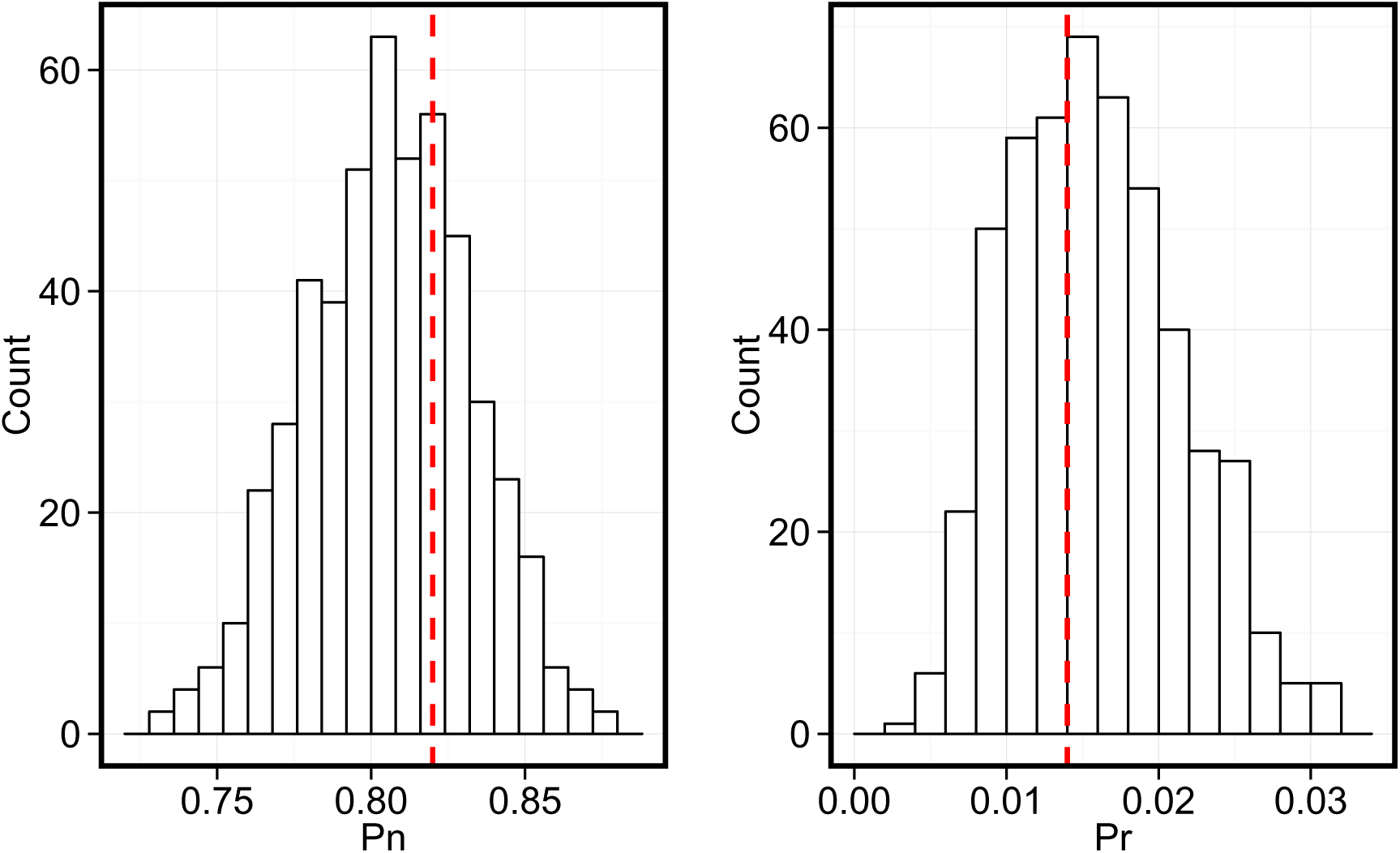
Example of PLS estimation using simulated network. The distributions of predicted *p_n_* (left) and *p_r_* (right) values for 500 networks simulated using *p_n_* = 0.82, *p_r_* = 0.014 are plotted, along with the real values (red dashed line).

**SI Figure 4:**
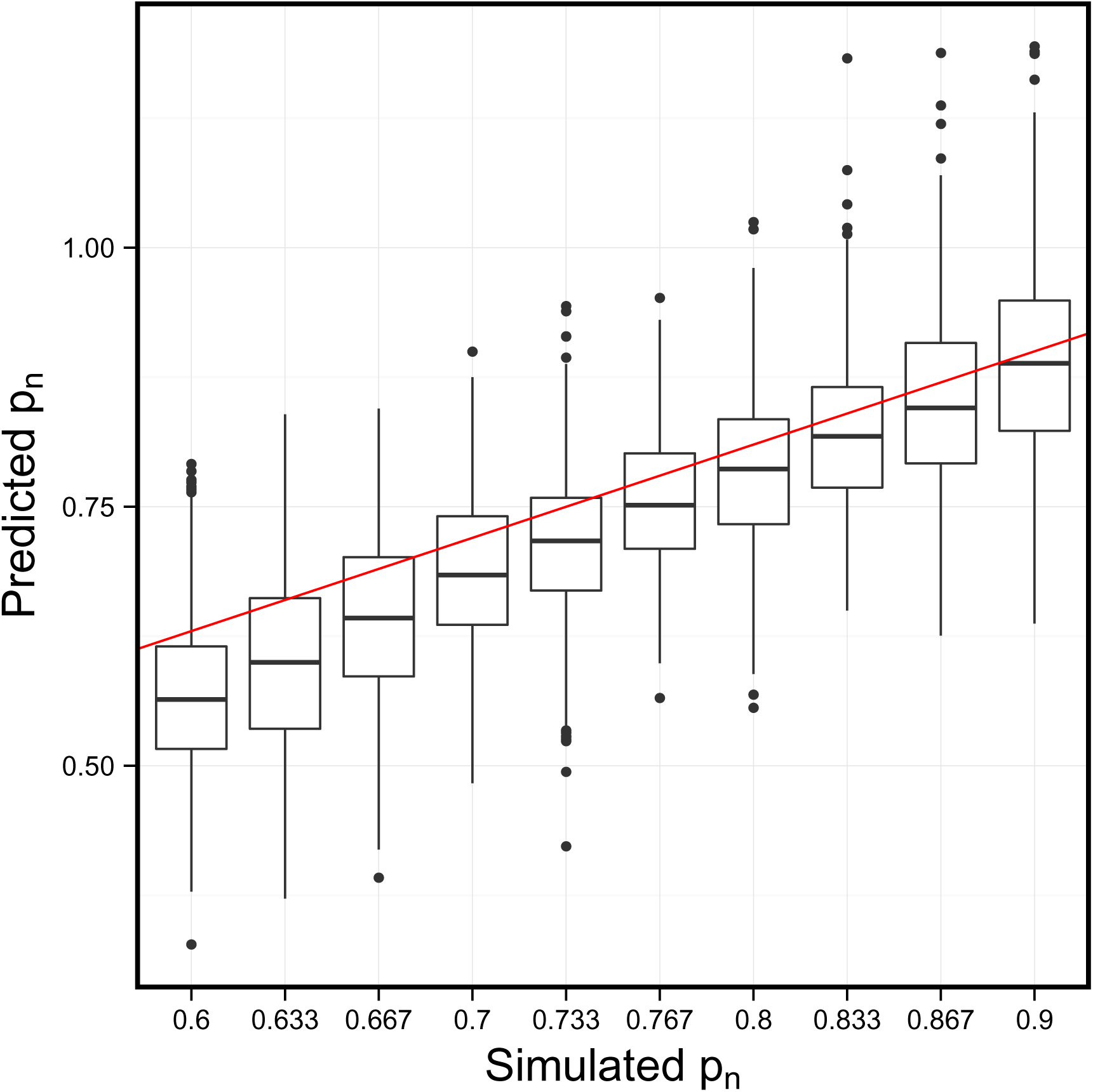
The PLS method estimates parameters with reasonable accuracy. Distributions of predicted *p_n_* values (box plot), compared to simulated values (red line). The predictions were generated after fitting a PLS regression to degrees and clustering coefficients of simulated networks. Thus, it is possible to predict the model parameter values when given an observed network.

**SI Figure 5:**
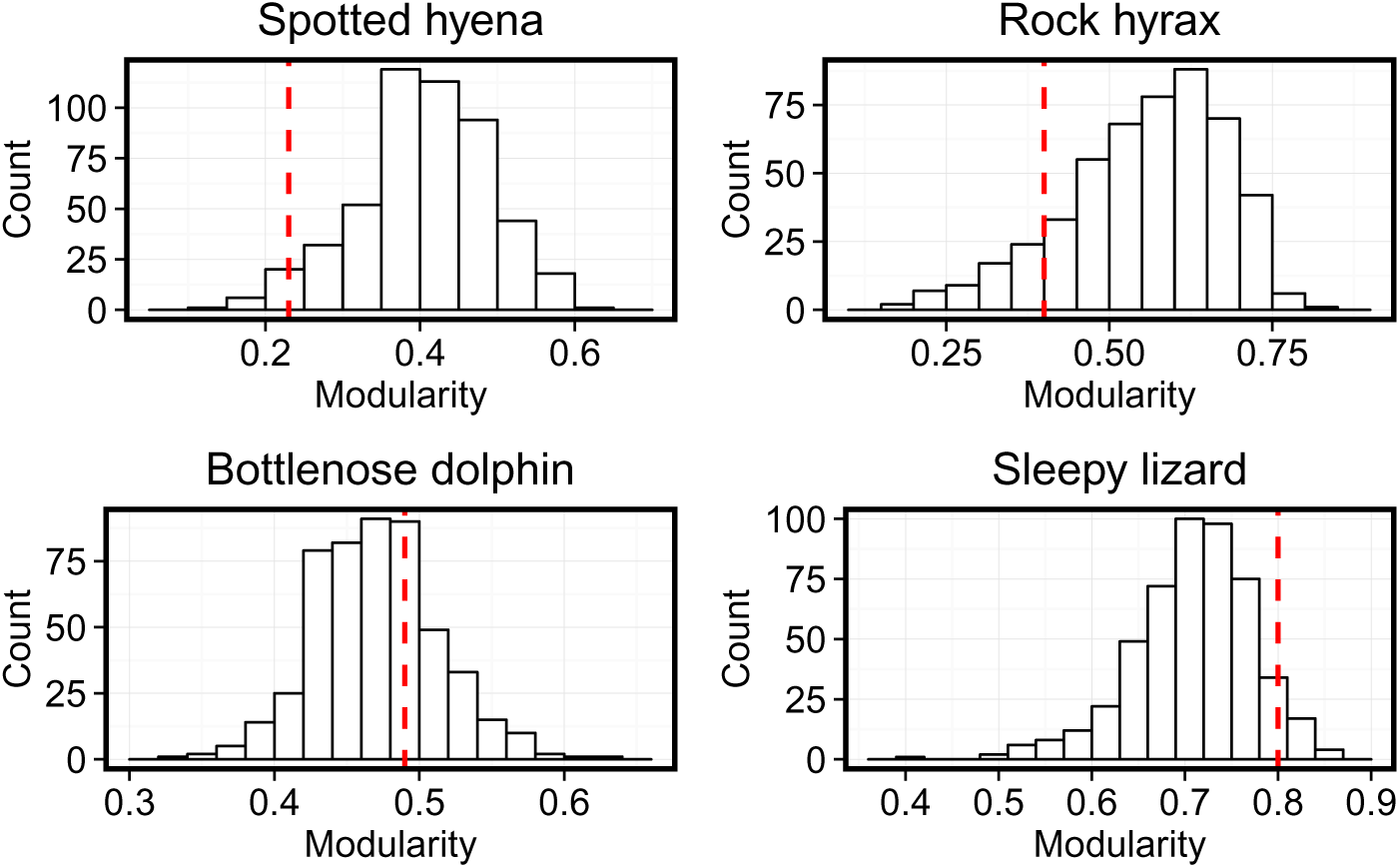
Modularity of networks with fitted model parameters. The network modularity of simulated networks from our model (distribution), compared to modularity of observed networks (red line). Modularity was calculated after partitioning the network to communities using the Walktrap algorithm. In all four species, the observed modularity could be generated by the model, i.e. was not an outlier.

**Table 1:** Fitted parameter values. Predicted parameter values used in the simulations for each species (predicted by partial least squares (PLS) regression; and predicted using analytical approximation of the mean degree and clustering coefficients). PLS values are used in Figure **3**. SI figure S2 plots the same using the estimates from the analytical approximations. A more detailed description of the PLS regression procedure and analytic approximation is in the Methods.

| Species | PLS |  | analytical |  |
| --- | --- | --- | --- | --- |
| | $p_n$ | $p_r$ | $p_n$ | $p_r$ |
| Spotted hyena<sup>13</sup> | 0.85 | 0.017 | 0.83 | 0.018 |
| Rock hyrax<sup>40</sup> | 0.75 | 0.007 | 0.66 | 0.026 |
| Bottlenose dolphin<sup>39</sup> | 0.50 | 0.028 | 0.43 | 0.036 |
| Sleepy lizard<sup>42</sup> | 0.51 | 0.007 | 0.38 | 0.012 |

Next, we tested if social inheritance can result in heritability of indirect network traits in social networks. Direct network traits (individual network traits that depend only on direct association with others, i.e. on the immediate social environment), such as degree, will by definition be heritable when *p_n_* is high and *p_r_* low. To see if this also holds for indirect network traits (traits that may depend also on associations between other individuals), we measured the correlation between parent and offspring betweenness centrality (which quantifies the number of times a node acts as a bridge along the shortest path between two other nodes; see Methods) for a set of social inheritance (*p_n_*) values. As Fig. **4** shows, high probabilities of social inheritance results in a pattern of heritability. In other words, when individuals are likely to copy their parents in forming social associations, the resulting network will exhibit heritability of centrality traits, although the only heritability programmed into the model is that of social inheritance and stochastic bonding. Similar patterns obtain for local clustering coefficient and eigenvalue centrality (Figures S**7** and S**8**).

**Figure 4:**
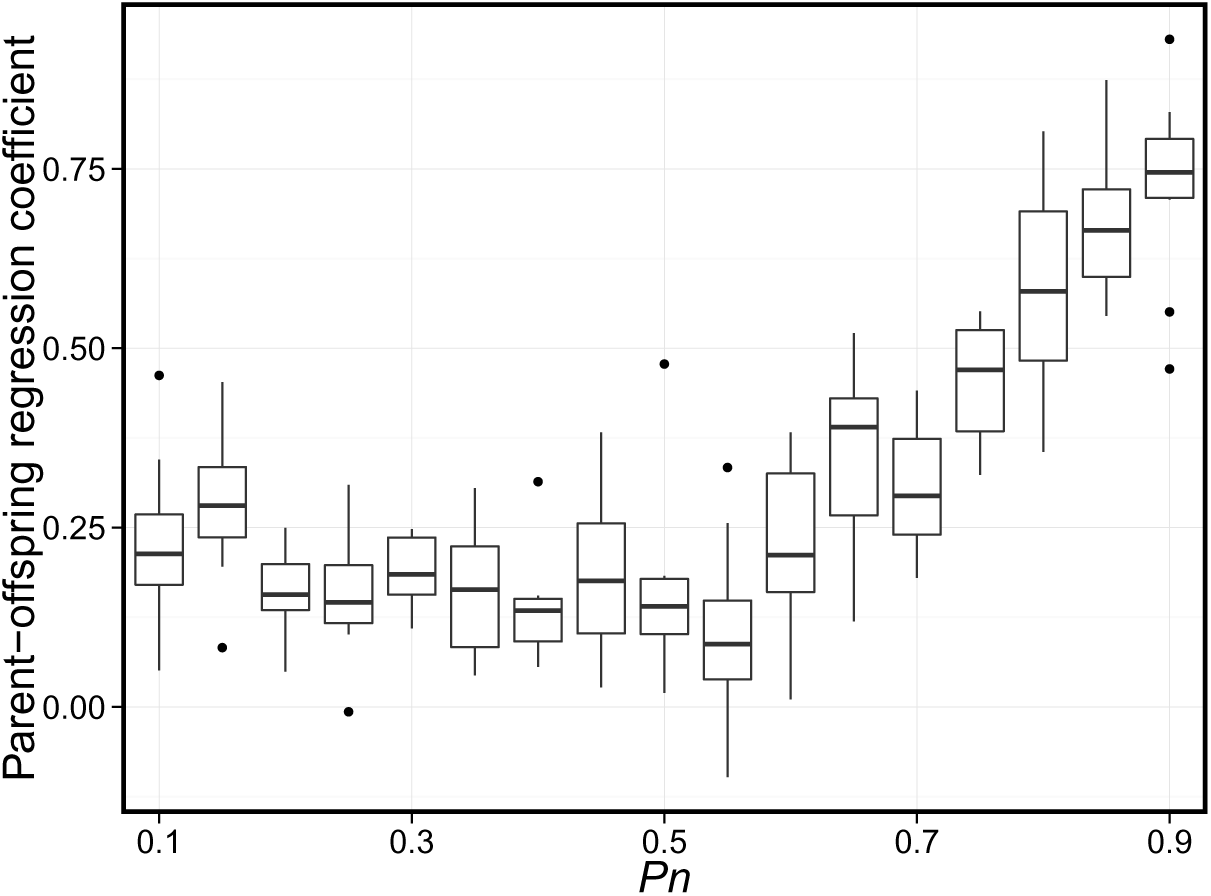
Social inheritance leads to the appearance of genetic heritability of network traits. The regression of betweenness centrality among parents and their offspring as a function of the strength of social inheritance (*p_n_*). The bottom and top of the box mark the first and third quartiles. The upper whisker extends from the hinge to the highest value that is within 1.5*IQR of the hinge, where IQR is the interquartile range, or distance between the first and third quartiles. The lower whisker extends from the hinge to the lowest value within 1.5*IQR of the hinge. Data beyond the end of the whiskers are outliers and plotted as points. Ten replications were run for each *p_n_* value. Parameter values: simulation steps=2000, *N* = 100, *p_r_* = 0.01.

**SI Figure 7:**
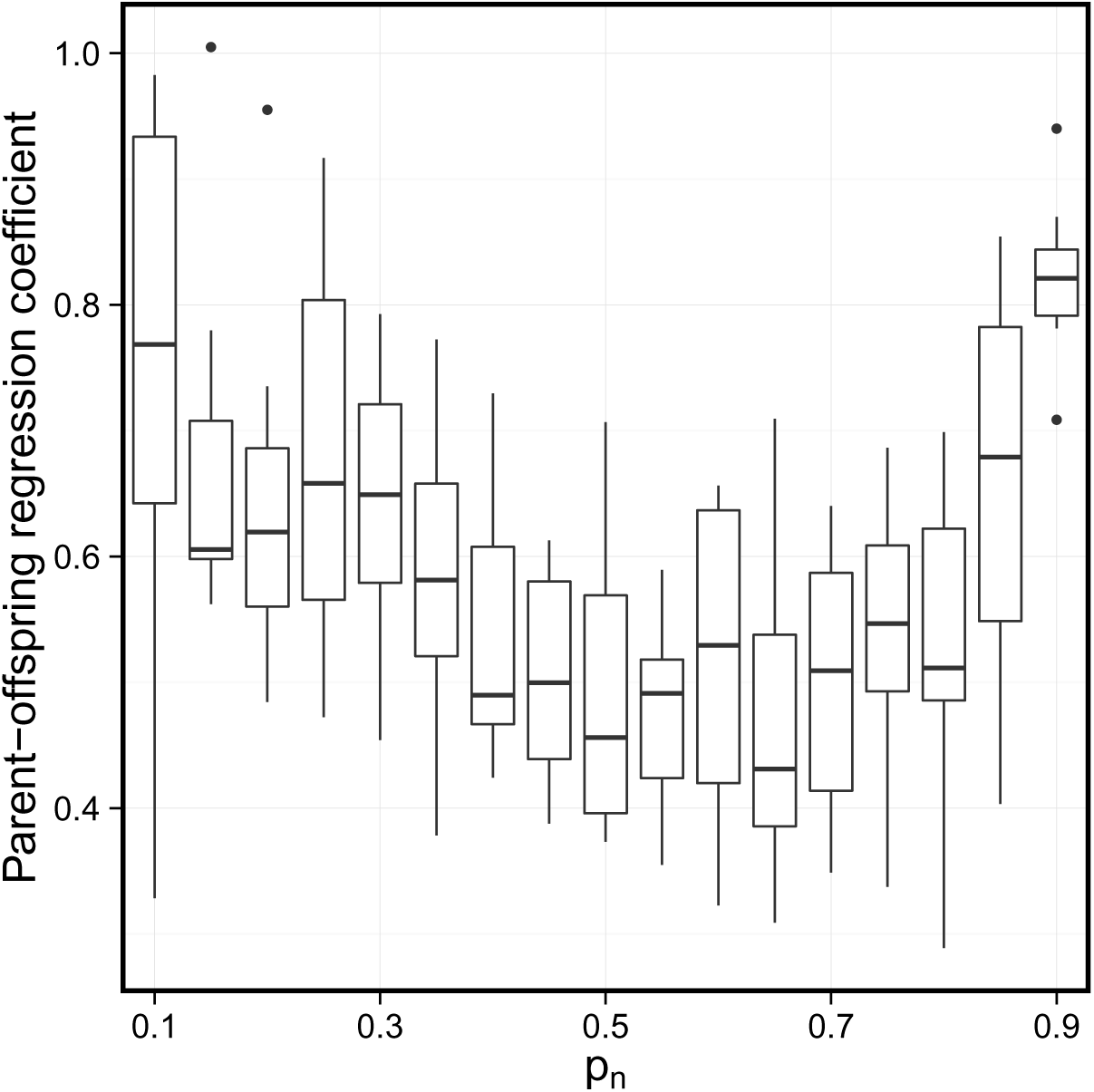
The heritability of local clustering coefficients. The figure depicts the regression of clustering coefficients among parents and their offspring as a function of the strength of social inheritance (*p_n_*). The bottom and top of the box mark the first and third quartiles. The upper whisker extends from the hinge to the highest value that is within 1.5*IQR of the hinge, where IQR is the inter-quartile range, or distance between the first and third quartiles. The lower whisker extends from the hinge to the lowest value within 1.5*IQR of the hinge. Data beyond the end of the whiskers are outliers and plotted as points. Ten replicate simulations were run for each *p_n_* value. Parameter values: simulation steps=2000 (parent-offspring regression calculated for the last 100 offspring born), *N* =100, *p_r_* = 0.01.

**Figure 1:**
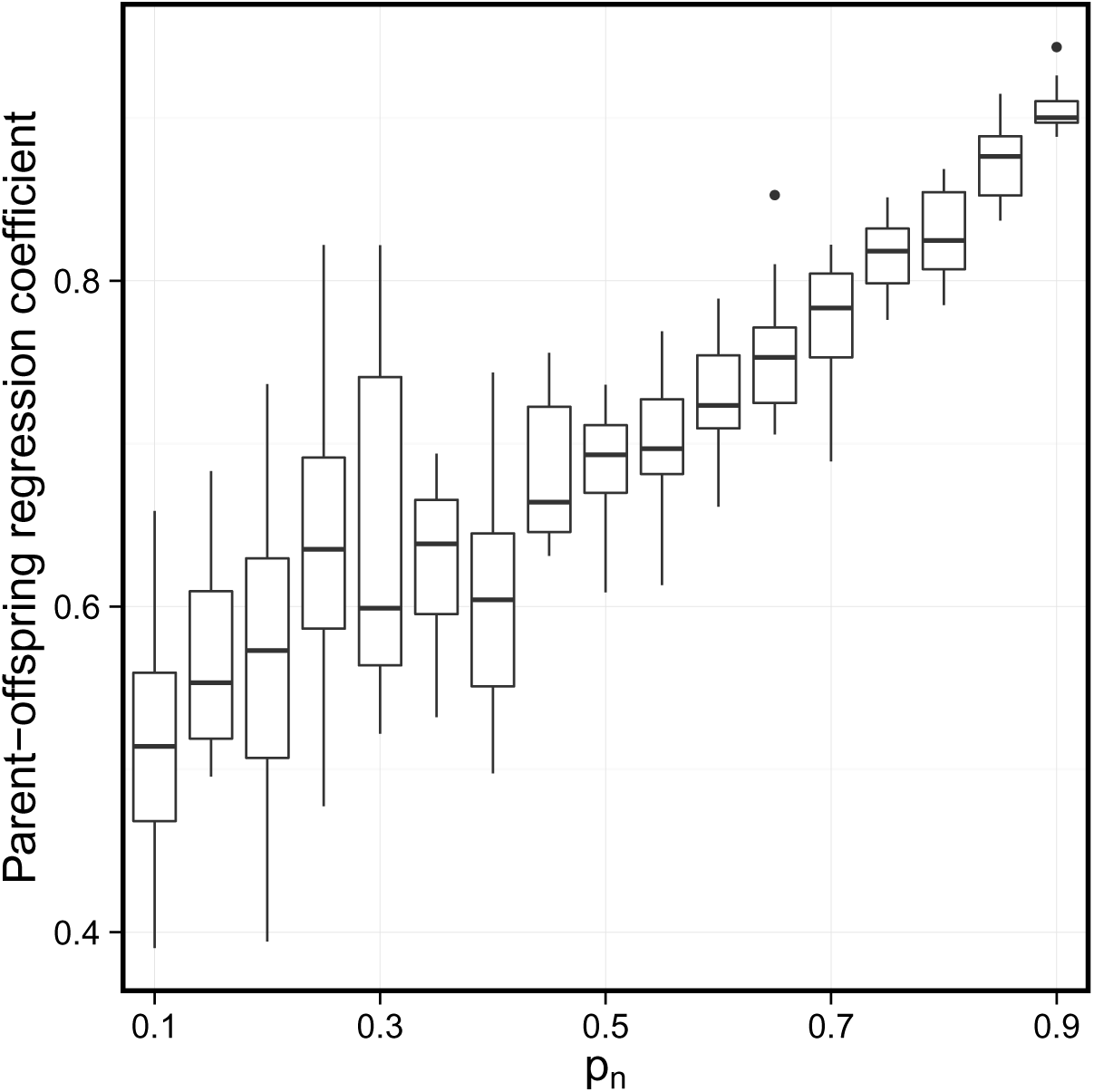
The heritability of eigenvector centrality. The figure depicts regression of eigenvector centrality among parents and their offspring as a function of the strength of social inheritance (*p_n_*). The bottom and top of the box mark the first and third quartiles. The upper whisker extends from the hinge to the highest value that is within 1.5*IQR of the hinge, where IQR is the inter-quartile range, or distance between the first and third quartiles. The lower whisker extends from the hinge to the lowest value within 1.5*IQR of the hinge. Data beyond the end of the whiskers are outliers and plotted as points. Ten replications were run for each *p_n_* value. Parameter values: simulation steps=2000 (parent-offspring regression calculated for the last 100 offspring born), *N* = 100, *p_r_* = 0.01.

Finally, we tested the effect of social inheritance on assortativity, i.e. the preference of individuals to bond with others with similar traits. We simulated networks where each individual had one trait with an arbitrary value between 0 and 1. Newborns inherited their mother’s trait with probability *1-μ*, where *μ* is the rate of large mutations. If a large mutation happened, the newborn had a random uniformly distributed trait value; otherwise, its trait was randomly picked from a Gaussian distribution around the mother’s trait, with variance σ^2^. Individuals followed the same rules of the basic model when forming social bonds. Hence, individuals did not explicitly prefer to bond with others with the same trait value. Nevertheless, the assortativity coefficient was significantly higher than in random networks, in which the trait values were re-assigned randomly (Figure **5**).

**Figure 5:**
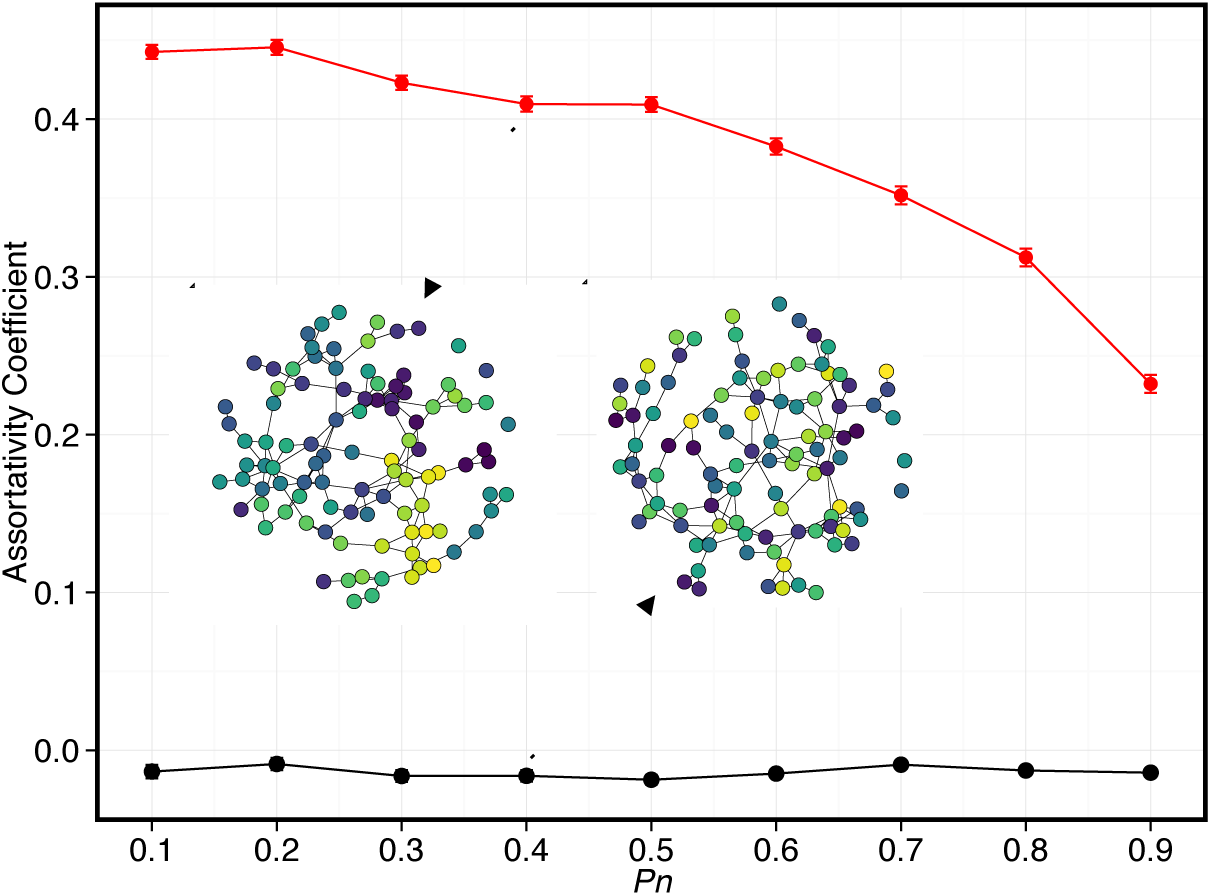
Assortativity as a consequence of social inheritance. Illustration of assortativity without explicit assorta ti ve preference. Dots and notches note assortativity coefficients and standard errors, respectively, for model networks (red), and shuffled networks, where trait values were reassigned randomly. Inset networks illustrate examples from the two groups. Circle colors represent arbitrary continuous trait values. Lines represent social bonds between individuals. Parameter values are the same as in Figure **4**, with mutation probability *μ* = 0.05

As an alternative model generating assortativity, we considered an explicit assortativity model, in which newborns explicitly prefer bonding with those with similar traits. Although this model unsurprisingly generated networks with high assortativity (mean assortativity coefficient±SEM: 0.53±0.006 compared to -0.01±0.002 in networks with randomly shuffled trait values), it failed to recover the high clustering and modularity observed in networks generated by social inheritance and in the data (Supporting Information, figures S**9** and S**10**). This result further suggests that assortativity might be a byproduct of social inheritance rather than a driving force of social network structure. A more generalized preferential attachment model, described in section SI 7, shows the converse is not true, i.e., that network patterns generated by social inheritance do not arise as a byproduct of genetically inherited traits and association preferences (see Discussion for more).

**SI Figure 9:**
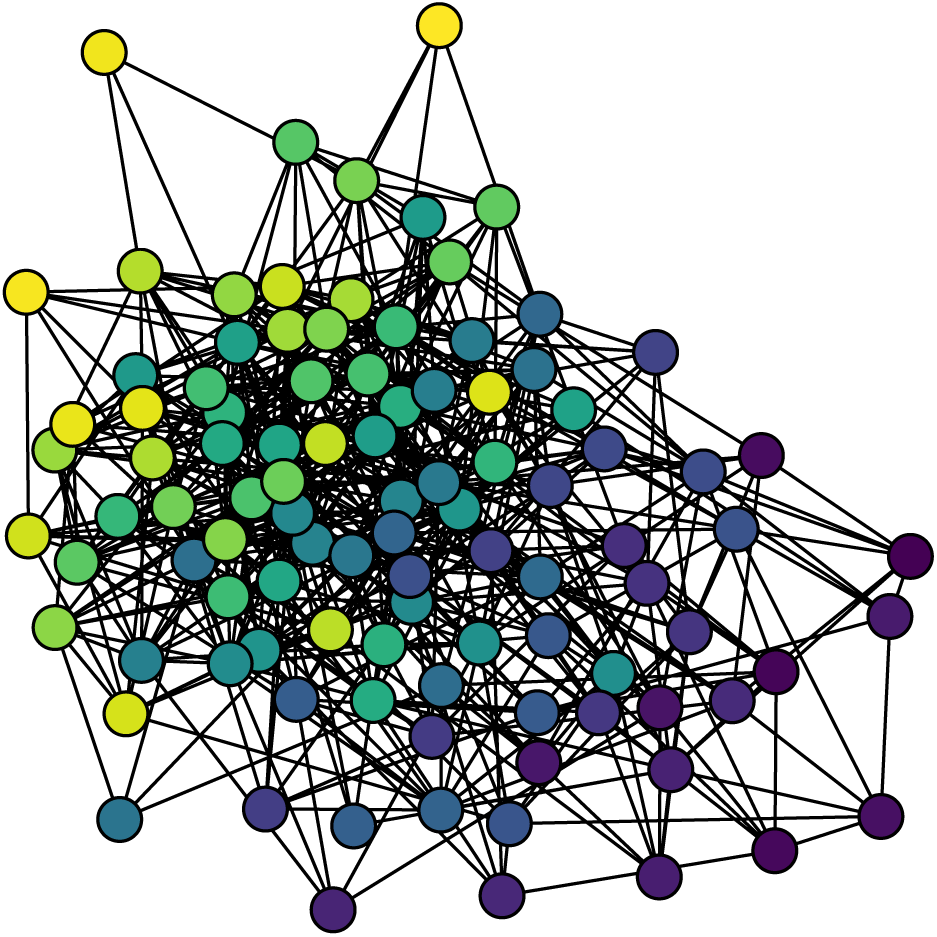
Example network generated with the assortative attachment model. An example of a social network resulting from the explicit assortativity model, in which newborns are more likely to connect with similar individuals. Colors represent the values of an arbitrary trait, considered when forming bonds. See text for model definition and simulation parameters.

**SI Figure 10:**
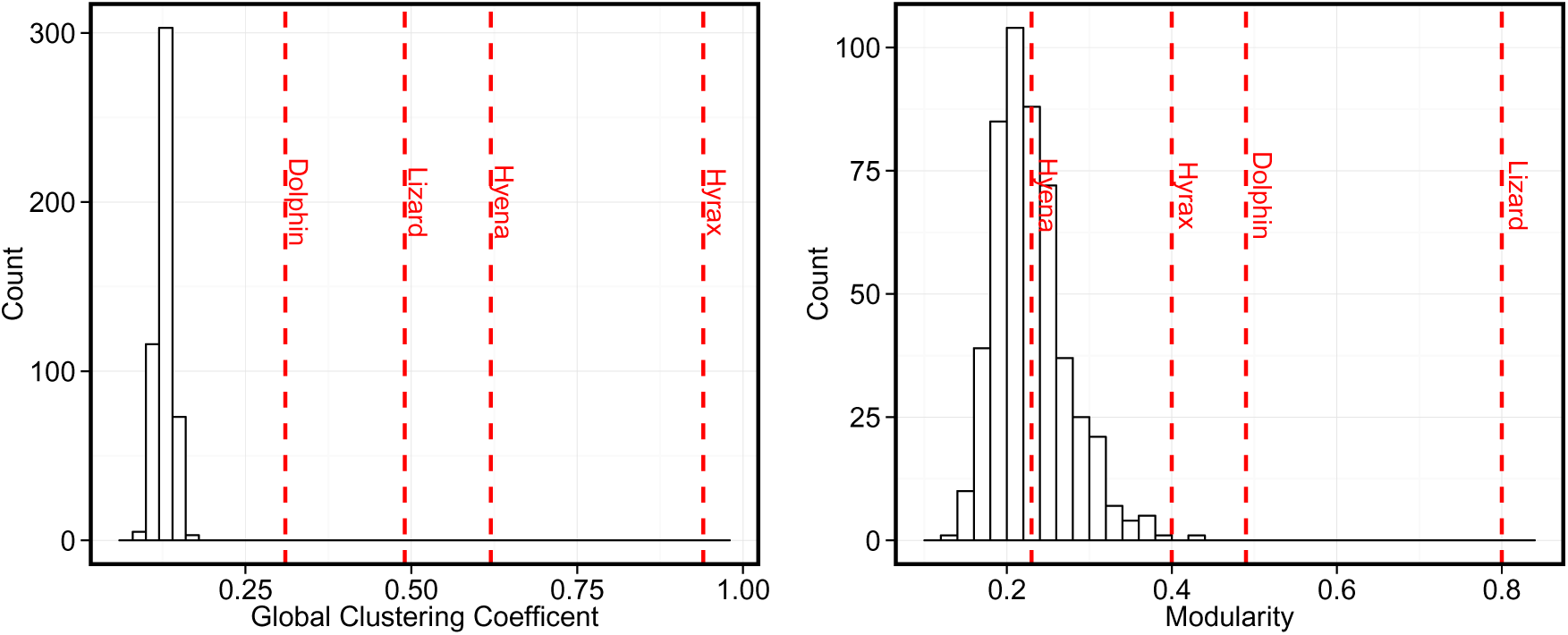
Clustering and modularity in the assortative attachment model. A comparison of the global clustering coefficient and modularity of 500 networks resulting from the explicit assortativity model (see text for details) to the values of observed networks of four species. Distributions show value of network measures for model networks. Red line show values for observed networks. The global clustering coefficient of all model networks is much lower than that of observed networks. Similarly, the modularity of model networks is lower than observed networks, except for the spotted hyena.

## Discussion

Our model provides a step towards a general theory of social structure in animals that is grounded in social and demographic processes. Our approach is to use dynamic generative models based on simple processes to predict network-scale patterns that those processes are expected to produce, and compare them to observed networks. Such an approach has been widely and productively used in network theory and social sciences^43,44,27^, as well as other subfields of ecology^31,32^ but not in animal social networks. Our work addresses this gap. Our main result is that the combination of neutral demography and social inheritance can replicate important properties of animal social networks in the wild.

In particular, we show that our model can capture essential features of social networks of four different species in the wild, including not just the degree distribution and modularity, but also the clustering coefficient distribution, in contrast to most studies of social network formation. Clustering is an important feature of social networks, distinguishing them from other types of networks, such as transportation networks and the internet^30^. Theory predicts that clustered networks are more conducive to cooperation^45^, and empirical studies document a tendency to close triads^40,13^, suggesting that it might be a generally adaptive feature of social structure. Nevertheless, many previous models of sociality and network formation fail to account for the high clustering observed. For example, whereas preferential attachment can reconstruct the degree distribution of social networks, it fails to capture their high degree of clustering^27^. The social inheritance process is crucial to the formation of cohesive clusters in social networks because it biases newly formed connections to those that close triads of relationships.

Social inheritance requires a behavioral mechanism that facilitates introduction of newborns to their mother’s social partners. As in many species young individuals tend to follow their mothers, it is easy to think about such a passive mechanism: young individuals are introduced to other individuals by spending time with their mother’s partners. This process is consistent with the long-held view that mother-offspring units are fundamental to social structure^46^. Direct evidence for social inheritance comes from Goldenberg et al.^36^, who documented the tendency of female African elephants to “inherit” the social bonds of their mothers, driving network resilience. Moreover, in many species group members show active interest in newborns^47^, promoting the initiation of a social bond between newborns and their mother’s partners. Further work can test if initial interest in newborns later translates to stronger social bonds with individuals reaching adulthood. We note that social inheritance does not necessarily require an active process of “introductions” but can also happen passively, for example as a result of spatial fidelity among group members. Our model is agnostic with regard to the mechanism of social inheritance. That being said, the fitted model parameters for the four networks vary in ways that are suggestive for socio-ecological factors: for hyenas and hyraxes, we find high *p_n_* values, which may reflect the strong philopatry in these societies. In contrast, the relatively low fitted value of *p_n_* in dolphins may reflect their multi-level society featuring mother-son avoidance^48^.

We make a number of simplifying assumptions, such as no individual heterogeneity, or age - or stage-structure in our demography. Models of this type have a long and distinguished history in ecology and evolution^49^, and in the same spirit, we do not believe that nature is actually as simple as we model it. Nonetheless, the fact that this very simple model (but not other simple models, e.g., see SI 6, SI 7) can reproduce important aspects of real networks suggests that the social inheritance of connections is likely to be important in structuring social networks. Even though the details will no doubt vary across species and contexts, this simple, quantifiable process can explain observed variation in social networks. For example, our model does not treat sex-specific dispersal, a mechanism that results in different social environments for the two sexes. Nevertheless, there is evidence for social bonding with familiar individuals after dispersal^50^. This suggests that even after dispersal, individuals may “inherit” the social bonds of certain conspecifics serving as role models. Another use of simple models such as ours is to serve as a base model to test the effect of additional factors. For instance, after fitting the model to an observed social network, one could test whether personality can explain the variance not explained by social inheritance and stochasticity. This can be attained by adding personality to the agent-based model as a factor that influences individual bonding decisions.

Our model has implications for how the inheritance of positions in social networks, which has important implications for social dynamics, is to be interpreted. For example, Fowler et al. ^51^ found that in humans, network traits such as degree and transitivity were heritable. In rhesus macaques, Brent et al.^52^ found that indirect network traits such as betweenness are more heritable than direct ones. In contrast, a study of yellow-bellied marmots, *Marmota flaviventris*, presented evidence for heritability of social network measures based on direct interactions^53^, but not indirect interactions. Taken together, these studies show that network position can be heritable, but have not been able to elucidate the mechanism of inheritance. It is not unlikely that some social network traits are genetically inherited; for example, individuals might genetically inherit social preferences from their parents that lead them to connect to the same individuals. In SI 7 we show that such a mechanism is unlikely to account for the observed levels of clustering. Therefore, our work suggests that at least some of the heritability of network traits might be social (as opposed to genetic), from individuals copying their parents. This prediction is borne out by recent studies in elephants^36^. Importantly, while these previous studies attempt to control for effects of the social environment at the group or lineage level using quantitative genetics methods e.g.^54^, they were not designed to distinguish social inheritance at the individual level from genetic inheritance. Studying the dynamics of social bond acquisition can be a way to separate genetic and social inheritance.

Another robust finding in network science and animal behavior is that individuals tend to connect to others with traits similar to themselves (e.g.,^55,56,57^). This assortativity is crucial for social evolutionary theory, as the costs and benefits of social interactions depend on partner phenotypes. Recent work has found that assortative mating can arise without assortative preferences, as a result of dynamic processes in a closed system^58^. Our results show that social inheritance can lead to high assortativity in the absence of explicitly assortative preferences for social bonding. Indeed, an alternative model based on explicit assortativity failed to reconstruct topological features of observed networks. Empirically, our results call for a careful assessment of networks with apparent phenotypic assortment, and controlling for social inheritance. This will be difficult to do with only static network data, but will be feasible for species with long-term data on the network dynamics.

Our work points to several interesting avenues to be explored in future research. First, we used binary networks to describe the strength of social bonds that are inherently on a continuous scale^11,59^. Weighted networks that can describe the delicate differences in the strength of social bonds between individuals would be more relevant in some cases. Future generative models can consider varying bond strength by coupling a weighted network model with a model of behavioral dynamics of social bond formation for pairs of individuals. Second, even though our model is extremely simplistic, most of its mathematical properties (including probability distributions over network measures such as the degree distribution) are analytically intractable, which makes model-fitting a challenge. Methods such as approximate bayesian computation^60^, coupled with dimensionality reduction techniques^61^ can be used to develop algorithms for estimating parameters of the model and also incorporate more information about individual variation and environmental effects (See SI 2 for more). Additionally, long-term datasets on social network dynamics can allow estimation of the social inheritance and random bonding parameters *p_n_* and *p_r_* directly. Lastly, our model does not consider changes in social bonds after these were established. Although this assumption is supported by empirical findings concerning bond stability in some species^13,12^, future models in which this assumption is relaxed should be developed. We also assume a single type of bond between individuals, whereas in nature, different social networks exist for different kinds of interactions (e.g., affiliative, agonistic, etc.). Such “multilayer networks”^62^ represent an important future direction.

In conclusion, the theory we present here is based on the idea that social networks should be regarded and analyzed as the result of a dynamic process^63^ that depend on environmental, individual, and structural effects^13^. Our work represents a first step in developing a theory for the structure of social networks and highlights the potential of generative models of social and demographic processes in reaching this goal.

## Methods

### Expected mean degree and clustering coefficient

In this section and the next, we characterize some important aspects of our model analytically. First, we can write a simple approximation of the expected mean degree, *d̅*, of a network changing according to our model at stationarity. To do that, we note that at stationarity, killing an individual at random is expected to remove *d̅* connections from the network. After this individual is removed, the average degree of the network becomes: 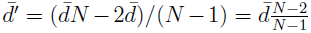. The expected degree of the connections made by the newborn is then: *p_b_* + *d̅p_n_* + (*N* – 2 – *d̅*′)*p_r_*. At stationarity, the links destroyed and added need to be the same on average, so we can write:

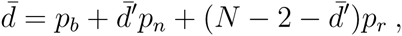

and solve for *d̅* to obtain:

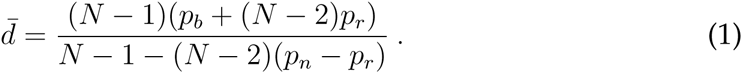

This approximation gives an excellent fit to simulated networks across all ranges of mean degree (Figure **2**).

We can also approximate the expected mean clustering coefficient of a network at stationarity using a similar stationarity argument (see SI 1 for the derivation). Our simulations (Figure **2**) show this approximation is very good except for the combination of very low *p_r_* and low to moderate *p_n_*, where it significantly overpredicts clustering.

Using the approximations for the mean degree and clustering coefficients, assuming *p_b_* = 1, and taking *N* to be the observed network size, we can estimate the *p_n_* and *p_r_* values for an observed network. In simulated networks, this approach generally yields accurate predictions except for the combinations of high *p_n_* and high *p_r_* (where it underestimates *p_r_*) and low *p_n_* and very low *p_r_* (where it overestimates *p_r_*). Our estimates of *p_n_* and *p_r_* for the four empirical networks from the analytical approximation are given in Table 1.

### Expected degree distribution

Finally, we characterize the expected degree distribution in our networks using a mean-field model. We denote the degree distribution by *ϕ_d_* for 0 ≤ *d* ≤ *N* – 1. In other words, *ϕ_d_* is the probability that a randomly selected individual in the population has degree *d*.

Consider a focal individual that has degree *d* at time period *t*. In period *t*+1, the probability that this individual increases its degree by one, 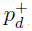, is:

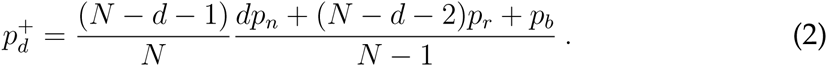

The first fraction in (2) is the probability that the individual selected to die is not connected to the focal individual, while the second fraction is the expected probability that the newborn individual born to one of the remaining *N* – 1 individuals becomes connected to the focal individual.

The probability of a focal individual’s degree *d* (> 0) going down by one, 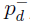 is likewise given by

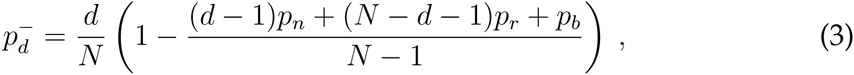

which is simply the probability that the individual selected to die is connected to the focal individual, multiplied by the probability that the newborn individual does not connect to the focal individual.

Next, we need the probability that a newborn is born with *d* connections, denoted by *b_d_*. To compute this probability, we assume *p_b_* = 1 (the extension to *p_b_* < 1 is trivial), so that the newborn always connects to its parent, then *b_d_*(*ϕ*) is given by (for *d* ≥ 1; *b*_0_ = 0 in that case):

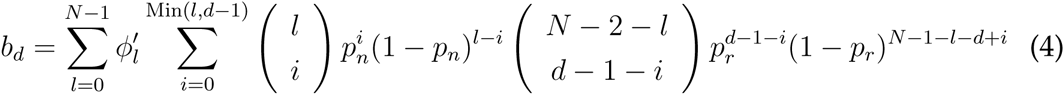

where the inner sum is the probability that an offspring of a parent of degree *l* is born with degree *d*, and the outer sum takes the expectation over 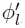, the expected degree distribution after the death of a random individual, which for 0 ≤ *l* ≤ *N* – 1 is given by:

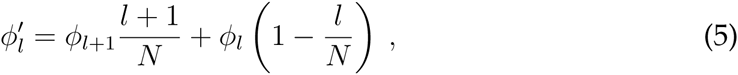

reflecting the facts that the death of a random individual does not change the expected frequency of individuals that had degree *d before* the death, but with each death, an individual with degree *d* has a probability *d*/*N* of becoming degree *d* – 1.

Putting everything together, we can write the rate equation for the mean-field dynamics of the degree distribution^28^:

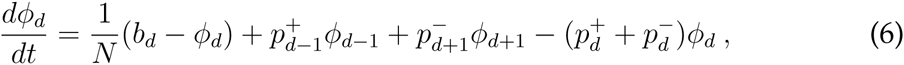

where the first termis the rate of change in the frequency of degree *d* caused by the replacement of individuals of degree *d* by death and birth, and the rest of the terms give rates of degree changes due to losing and gaining connections.

Setting equation (6) equal to zero for all *d* and solving the resulting *N* equations, we can obtain the stationary degree distribution. We were unable to obtain closed-form solutions to the stationary distribution, but numerical solutions display excellent agreement with simulation results (see Figure **3**). It is worth noting that although the 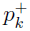 and 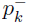 terms are similar to models of preferential attachment with constant network size e.g.^28^, these models assume that each new addition to the network has exactly the same degree, whereas in our model, the number of links of a newborn is distributed according to equation (4). Furthermore, the degree distribution does not capture the clustering behavior of preferential attachment models, which generate much less clustering than our model for a similar mean degree (results not shown), consistent with results in growing networks^27^.

### Simulation process

We initialized networks as random graphs, and ran them long enough to converge to steady state, which we evaluated by the mean degree distribution of ensembles matching the expected degree distribution, mean degree and clustering values derived analytically. The time to convergence to steady state depends on the network size, *p_n_*, and *p_r_*: we found as a rule of thumb that 10 times the network size (i.e., 10 complete population turnovers on average) is enough for networks to come to stationarity, hence our choosing of 2000 steps for network size of 100. The only exception is with *p_n_* close to 1 (and to a lesser extent, *p_r_* very close to zero), where we find that convergence can take significantly longer.

### Fitting models to observed networks

To obtain estimates of parameter values *p_n_* and *p_r_* from observed networks, we used two methods: (i) a computational approach using dimensionality reduction on the degree and local clustering distributions of simulated networks, and (ii) an analytical approach using approximations of the mean degree and local clustering coefficients. In this subsection, we describe the dimensionality reduction approach. For each empirically observed network, we ran the model with 10000 random values of *p_n_* and *p_r_* between 0 and 1, and the network size was set to match the observed network. We then used partial least squares regression, using the R package *pls* (version 2.4-3), to obtain a regression of the network degree and clustering coefficient distributions on *p_n_* and *p_r_*. Based on the regression formula, we predicted the values of *p_n_* and *p_r_*. The values predicted by the regression were sufficient to simulate networks that were close in their degree and clustering coefficient distributions to the observed networks. The values given in Table 1 are the result of the PLS fit. They are meant to demonstrate the ability of the model to generate realistic looking networks. In the SI, we provide a verification of the method’s ability to obtain the values of *p_n_* and *p_r_*.

## Data

We compared the output of our model with observed animal social networks of four different species. For this analysis we used data from published studies of spotted hyena *(Crocuta crocuta*^13^), rock hyrax *(Procavia capensis*^40^), bottlenose dolphin *(Tursiops* spp.^41^), and sleepy lizard *(Tiliqua rugosa*^42^).

The hyena social network was obtained from one of the binary networks analyzed by^13^, where details on social network construction can be found. Briefly, the network is derived from association indexes based on social proximity in a spotted hyena clan in Maasai Mara Natural Reserve, Kenya, over one full year (1997). The binary network was created using a threshold retaining only the upper quartile of the association index values. Similarly, the hyrax network was described by^40^, and is based on affiliative interactions in a rock hyrax population in the Ein Gedi Nature Reserve, Israel, during a five-months field season (2009). The same upper quartile threshold on the association indices was used to generate a binary network. The dolphin network was published in^41^, and is based on spatial proximity of bottlenose dolphins observed over 12 months in Doubtful Sound, Fiordland, New Zealand. “Preferred companionships” in the dolphin network represent associations that were more likely than by chance, after comparing the observed association index to that in 20000 permutations. The lizard social network was published by^42^, and is also based on spatial proximity, measured using GPS collars. To get a binary network, we filtered this network to retain only social bonds with association index above the 75% quartile.

### Network measures

To study the networks produced by our model and compare them to observed networks, we used a number of commonly used network measures. Network density is defined as 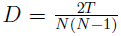 where T is the number of ties (edges) and N the number of nodes. The global clustering coefficient is based on triplets of nodes. A triplet includes three nodes that are connected by either two (open triplet) or three (closed triplet) undirected ties. Measuring the clustering in the whole network, the global clustering coefficient is defined as

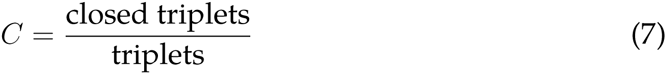

The local clustering coefficient measures the clustering of each node:

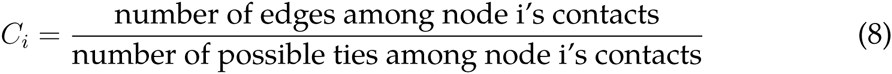

The betweenness centrality of a node *v* is given by

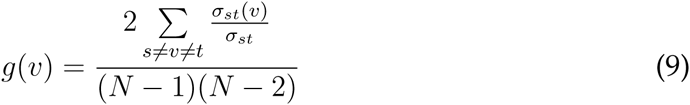

where *σ_st_* is the total number of shortest paths from node *s* to node *t* and *σ_st_*(*v*) is the number of those paths that pass through *v*.

We detected network modules (also known as communities or groups) using the walktrap community detection method^64^. We used the maximal network modularity across all partitions for a given network. The modularity measures the strength of a division of the network into modules. The modularity of a given partition to *c* modules in an undirected network is

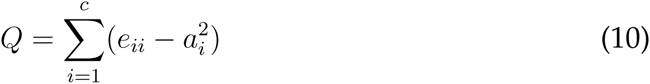

where *e_ii_* is the fraction of edges connecting nodes inside module *i*, and 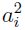 is the fraction of edges with at least one edge in module *i*.

Finally, we used the assortativity coefficient to measure how likely are individuals to be connected to those with a similar trait value^65^. For an undirected network, this coefficient is given by:

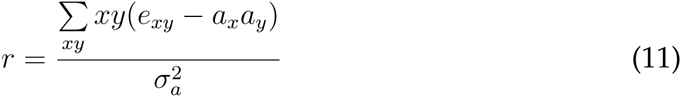

where *e_xy_* is the fraction of all edges in the network that connect nodes with traits *x* and *y, a_x_* is defined as 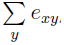, and 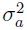 is the variance of the distribution *a_x_*.

## Acknowledgments

We are thankful to Kay E. Holekamp and Eli Geffen for sharing data, and to Robert Seyfarth, Elliot Aguilar, Çağlar Akçay, Jeremy Van Cleve, Slimane Dridi, and Tim Linksvayer for valuable comments. Comments from three anonymous reviewers helped improve the paper and are gratefully acknowledged. This study was supported by the University of Pennsylvania and NSF Grant EF-1137894.

## Supplementary Information

For “Social inheritance can explain the structure of animal social networks”

Amiyaal Ilany and Erol Akçay

### SI 1 Approximation for mean local clustering coefficient

Similar to the mean degree, we use a stationarity argument to calculate an approximation for the mean local clustering degree of a network by equating the expected clustering coefficient of a randomly killed individual with the expected change in the clustering coefficients of all remaining individuals with the birth of the newborn plus the expected clustering of the newborn itself:

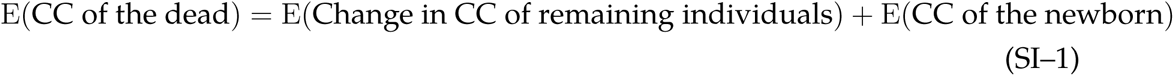

The expected clustering coefficient of an individual randomly selected to die is equal to c, the mean clustering coefficient. When an individual is killed, the clustering coefficient of its connections will in principle change, but one can show that the “typical” connection (i.e., one with degree *d̄* and clustering *c̄*) will not experience a change in its clustering coefficient. This can be seen by calculating the new clustering coefficient after death,

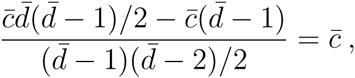

where the first term in the numerator is the expected number of closed triangles a typical connection of the dead individual had before, the second term the number of triangles that were removed by death, and the denominator is the number of all potential triangles after death.

The birth of a new individual changes the total of the clustering coefficients in two ways: (i) by changing the clustering coefficients of individuals connected to the newborn, and (ii) by adding the newborn with the newborn’s clustering coefficient. Let us calculate the first effect: the clustering coefficient of an individual with initial degree *d* and clustering coefficient *c* that becomes connected to the newborn is going to change as follows:

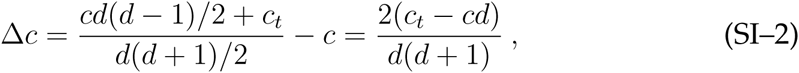

where the first term in the numerator of the middle part is the number of closed triangles amongst the focal individual’s connections before getting connected to the newborn, and *c_t_* is the expected number of closed triangles amongst the focal individual’s connections established by the newborn. The denominator is the total number of triangles after the focal individual gets connected to the newborn. To calculate *c_t_*, we need to consider the three kinds of connections of the newborns separately: its parent (with probability *p_b_*), its parent’s connections (with probability *p_n_*), and individuals not connected to its parent (with probability *p_r_*).

For the parent, the expected number of closed triangles generated by the newborn is simply

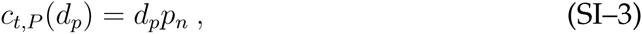

where *d_p_* is the degree of the parent. For a parent’s connection, each has on average *c_p_*(*d_p_* – 1) connections to other connections of the parent, which in turn have a probability of *p_n_* of getting connected to the newborn. Further, on average parent’s connections will have *d̄*̂/(*N* – 1)(*N* – *d_p_* – 2) connections to non-connections of the parent (where 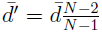, the expected degree of individuals after a death occurs), each of which have probability *p_r_* of getting connected to the newborn. Thus, for parent’s connections, we have

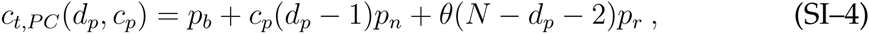

where *θ* = (*d̄*̂ – *c_b_*(*d_p_* – 1) – 1)/(*N* – *d_p_* – 2) is the probability a given non-connection of the parent is connected to a parent’s connection.

By a similar argument, one can write for non-connections of the parent:

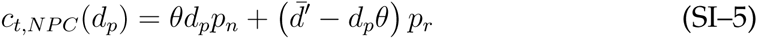

Thus, substituting *c_t_*,*P, c_t_*,*PC*, and *c_t_*,*NPC* into equation (SI-2), we can write for the expected total change in the clustering coefficient of existing individuals with the birth of the newborn, when the parent has degree *d_p_*:

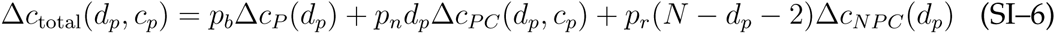

Next we need to calculate the expected clustering coefficient of the newborn, given the parent’s degree *d_p_* and clustering coefficient *c_p_*: *E*(*c*_NB_ |*d_p_*, *c_p_*). This number is the ratio of two random variables: *T*_*c*,_ the number of closed triangles that have the newborn as a vertex and *T*_*t*,_ the total number of pairs connected to the newborn, i.e.,

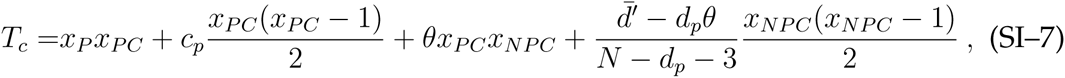

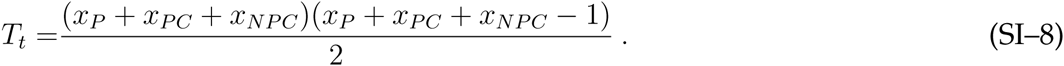

Here, *x*_•_ denotes the number of connections of the newborn to each class of individual *(P* for parent, *PC* for parent’s connections, and *NPC* for individuals not connected to the parent). Thus, *x_P_* is distributed according to a Bernoulli distribution with probability *p_b_*, *x_PC_* a binomial with parameters *d_p_* and *p_n_*, and *x_NPC_* a binomial with parameters *N* – *d_p_* – 2 and *p_r_*. The fractions in the third and fourth term in *T_c_* give the expected density of connections between a parent’s connection and non-connection, and amongst the non-connections, respectively. The expectation of the ratio of two random variables *T_c_* and *T_t_* can be approximated by their moments as follows:

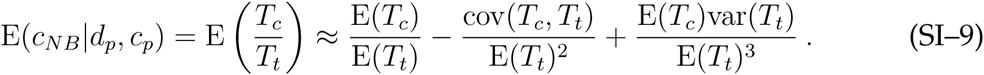

Using the distributions of *T_c_* and *T_t_*, computing (SI-9) is a straightforward if tedious calculation.

For the final step in our computation, we assume that the parent is chosen at random from the population, so has expected degree *d_p_* = *d̄*′, and clustering coefficient *c_p_* = *c̄*. Thus, our stationarity condition can be written as:

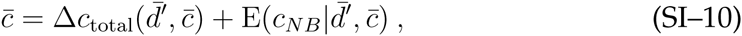

which can be solved for *c̄* analytically and *d̄* substituted from equation (1) to obtain an expression for *c̄* as a function of model parameters. We carried out our calculations in Mathematica 10 (Wolfram Research, Inc.). As Figure **2** in the main text shows, our approximations for the mean degree and clustering give an excellent fit to simulated networks, except for the mean clustering of networks with low *p_n_* and very low *p_r_*.

We can also use the equations (1) and (SI-10) to estimate the parameters *p_n_* and *p_r_* (assuming *p_b_* = 1) from the mean degree and clustering coefficient of a given network. In simulated networks, this method works well to estimate parameters (figure S**1**) except for high *p_n_* and moderately high *p_r_* values, where it tends to underestimate especially the *pr* values, and for low *p_n_* and very low *p_r_*, where it overestimates *p_r_*. Three of the four real-life networks we apply our model to fall comfortably in the region where the method yields reasonable accurate estimates (with *p_r_* values of the order of 0.01), with only the sleepy lizard network seemingly in a region where our estimate of *p_r_* somewhat inflated. Table 1 gives the values calculated for the four species, which produce networks that are similar to observed ones (figure S**2**) for hyenas, hyraxes and dolphins, but somewhat underpredicts clustering coefficients for the sleepy lizard network relative to the PLS method. The difference between the estimates for *p_r_* obtained from PLS and analytical approximation is consistent with the bias in the analytical estimators in simulated networks for low *p_n_* and *p_r_*.

### SI 2 Fitting the model to data: partial least square regression

Figure **3** shows that an objective procedure using partial least squares (PLS) regression can statistically identify values of *p_n_* and *p_r_* that will generate networks similar to the observed networks.

To verify the usage of PLS regression to fit our model to observed networks, we simulated networks using known parameter values and tested the predictions of PLS regression. Specifically, we simulated 10,000 networks from our basic model over 2000 time steps, using random *p_n_* and *p_r_* values. We then used PLS regression to fit the degrees and clustering coefficients to parameter values. We then simulated sets of 100 networks each using a given set of parameter values (p_n_ = 0.6 to 0.9, *p_r_* = 0.014) and checked whether the PLS regression fit could predict those values. For example, in SI Figure **3** we plot the distribution of predicted *p_n_* and *p_r_* values compared to the real values used to simulate the networks. SI Figure **4** shows the distribution of predictions for ten different values of *p_n_*, whereas *p_r_* was fixed at 0.014.

### SI 3 Modularity of model networks

Social networks feature higher modularity than random networks. That is, social networks can usually be partitioned into subgroups of individuals (communities in network jargon), more densely connected within than between those subgroups. To test another aspect of our model, we calculated the modularity of simulated networks after identifying the community (subgroup) structure. Modularity measures the strength of division into communities, where high modularity indicates dense connection between individuals within communities and sparse connections between individuals across communities. We used the Walktrap community finding algorithm, based on the idea that short random walks on a network tend to stay in the same community^64^. In all four tested networks (see main text), the modularity of the observed network was not an outlier in the distribution of modularity values of simulated networks. Thus, we could not reject the null hypothesis that the observed network belongs to the family of simulated networks, when considering their modularity (figure S**5**).

### SI 4 Two sex models

In the main text we presented the simplest model, in which the population was asexual. The basic model allows a newborn to choose any present individual as a role model to copy social associations. Here we show a version of the basic model for a sexual population. At birth, newborns are uniformly assigned a sex, and only females reproduce. Newborns copy only their mother’s associations. Thus males may form social associations when they are born, and also if a newborn connects to them, but they are not being copied by any newborn in terms of social associations.

**SI Table 1:** Parameter values used in the simulations of the two-sex model for each species in figure S**6**.

| Species | $p_n$ | $p_r$ |
| --- | --- | --- |
| Spotted hyena | 0.86 | 0.021 |
| Rock hyrax | 0.76 | 0.01 |
| Bottlenose dolphin | 0.50 | 0.033 |
| Sleepy lizard | 0.53 | 0.003 |

Fitting the two-sex model to data shows similar results to the basic model 1. This suggests that sexual reproduction is theoretically not a major determinant of social structure. Note that this does not mean that males and females play similar social roles in a population, but rather that if newborns tend to copy only one sex the resulting social structure is not very different.

We then tested two more models with sexual populations, in which the newborn may copy both parents with probability *p_n_*. In the first of these models, a newborn would copy any randomly chosen male and female as parents. In the second model, a newborn can be born only to connected pairs. Thus, in each iteration a pair of connected male-female was chosen as parents. Both these models generated networks that were not clustered, and could not be fit to observed data. This suggests that in natural populations individuals follow one role model, leading to the observed high levels of clustering. Theoretically, it is easy to see that if an individual follows multiple role models that is more similar to random connectivity, deviating from the structured observed networks of natural populations.

### SI 5 Heritability of social network traits

SI Figure **7** represents the parent-offspring regression for local clustering coefficient with varying *p_n_*. SI Figure **8** depicts the same for eigenvector centrality.

**SI Figure 6:**
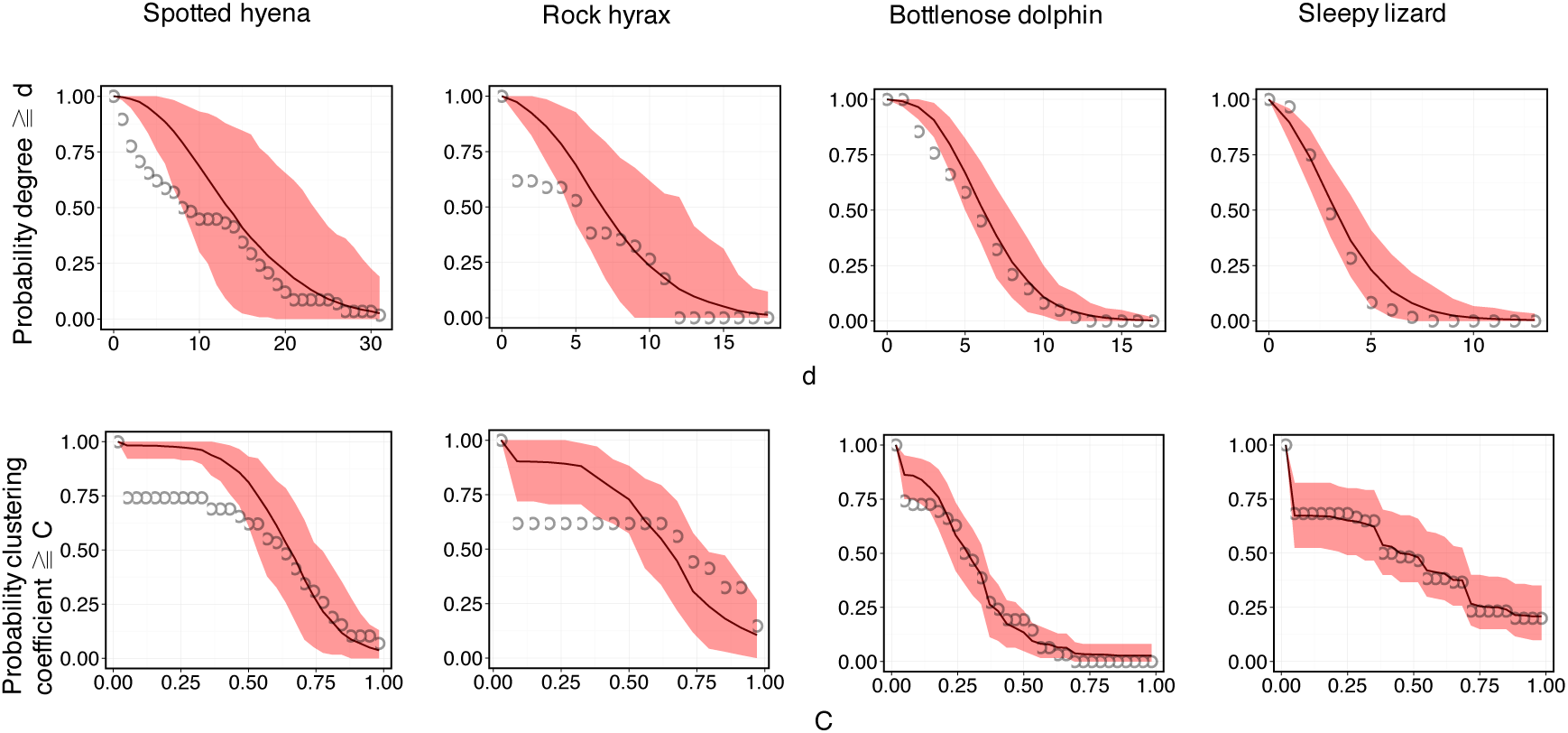
Comparing the two sex model output to networks of four species. Upper row: Cumulative degree distributions of observed and simulated networks. Lower row: Cumulative clustering coefficient distributions of observed and simulated networks. Black circles represent observed values. Red line notes mean values for 500 simulated networks (2000 simulation steps) with the same species-specific *p_n_* and *p_r_* values (given in Table 1), whereas light red area depicts 95% confidence intervals.

### SI 6 An alternative assortativity model

We constructed an alternative model of social network dynamics, focused on preference to form social bonds with other individuals with similar traits. The purpose of this model is to test the notion that explicit assortativity is the main factor determining network structure, as suggested empirically in various species. In this alternative model, newborns still bond their mother with probability *p_b_*, but then form bonds with all others with probability proportional to the similarity of an arbitrary trait value. The trait is inherited from the mother in the same manner as in the main model (see main text). Specifically, the probability of a newborn to connect with any other individual was defined as 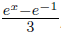, where *x* is the absolute difference in trait values of the newborn and a candidate individual. This term ensures the connection probability to be in a realistic range, resulting in networks with similar density to the mean density of the four observed networks (0.123, see main text).

Unsurprisingly, simulations of the explicit assortativity model (2000 time steps, 100 individuals, 500 replications) resulted in networks with high assortativity (figure S**9**). However, the resulting networks failed to reconstruct other important topological features of the observed networks, namely the global clustering coefficient and modularity (figure S**10**). The only exception was the spotted hyena, where modularity values, but not global clustering coefficient, matched modularity levels of the explicit assortativity model.

To conclude, a model of social structure where individuals base their social bonding almost exclusively on assortativity fails to reconstruct the topological features of observed networks in the tested species.

### SI 7 A generalized association preference model

A potential alternative interpretation of social inheritance is that it might arise as an epiphenomenon from genetically inherited association preferences (that may or may not be assortative): if individuals inherit their preferences for associating with certain types of individuals from their parents, they would be expected to be to associated with their parents’ connections more than unconnected individuals.

In this section we address this possibility by constructing a model to explore whether a more generalized model of co-inherited association preferences and traits might mimic the process of social inheritance. To generalize the assortative preferences model, we now assume each individual carries two traits, one describing a real-valued attribute (as in the assortment model above; we call this the “display trait”), and the other the preference for that trait (the “preference trait”). For example, if a focal individual has display and preference trait values (0.1,0.5), it is being preferred most by others with preference trait 0.1 but the focal individual prefers to associate with those having trait value 0.5. We assume both trait values are on a circle and normalize them to be between 0 and 1. We let both traits to be inherited from the parent when an individual is born, with (independent) deviations in each trait from parental values distributed according to *N*(0, *σ*). When an offspring *j* is born, it makes a connection to each existing individual *i* in the population with probability *e*^−*kd_ij_*^, where *k* is a positive constant and *d_ij_* is the shortest distance on the circle between the offspring *j*′s preference trait and the individual *i*′s display trait. Individuals are selected to die and give birth at random as in the basic model.

figure S**11** illustrates the results from this alternative model. It shows that although model parameters exist that generate realistic looking degree distributions, these generate networks that are far less clustered than the real-life networks. The reason is that when individuals connect to others purely based on their inherited display and preference traits, they tend to connect to both partners of their parents as well as other with similar traits that are not connected to their parents. The latter connections do not close triads, and hence the resulting network is much less clustered. Thus, purely genetic inheritance of association preferences (independent of parental connections) is insufficient to generate the process of social inheritance as a byproduct.

**SI Figure 11:**
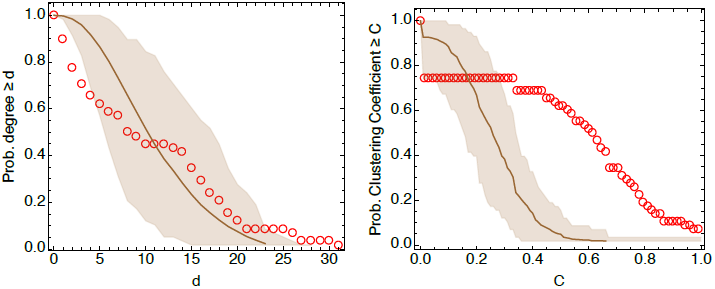
Generalized preference model does not produce enough clustering. Degree (left) and local clustering (right) distributions obtained from the generalized preference model described in section SI 7, compared with the spotted hyena network from 1997 (red open circles). The brown solid line gives the mean of the preferential association model with *N* = 58, *k* = 12 and *σ* = 0.05, and the shaded region the 10th and 90th percentiles in each panel.

### SI 8 The effect of varying network size

Population size might influence social structure in unknown ways. To test how changes in population size affect the resulting network, we simulated networks that grow or shrink in size. We then compared measures of the networks to those of stable networks, where the network size was kept constant. In a shrinking network model, we started the simulation with 200 individuals and ran it for the first 1000 time steps as a constant size network (one born and one dead at each time step). After 1000 steps we set the probability of each individual to die at any time step at 0.05, corresponding to an expected mortality of 10 individuals per time step initially. We kept the number of individuals born at each time step at one. We kept running the simulation until population size fell to 100 individuals, and compared network characteristics to a parallel simulation where the population size started out with *N* = 100 and held constant throughout. Similarly, in a growth model we started with 100 individuals for the first 1000 steps, and then changed the probability of each individual to die at a given time step to 0.001 (instead of 0.01 in a stable network size). We stopped the simulation when the network size increased to 200. Again, we compared these networks to networks that started out with *N* = 200 were kept constant throughout. We present results for a series of 15 parameter sets, where *p_n_* varied between 0.5 and 0.9 (5 values) and *p_r_* was one of 0.01,0.05, and 0.1. For each parameter set, we ran 100 replicate pairs of shrinking (or growing) and constant size networks. Figures S**12**, S**13**, and S**14** compare the network measures of stable to shrinking and growing networks, for the tested parameter sets.

**SI Figure 12:**
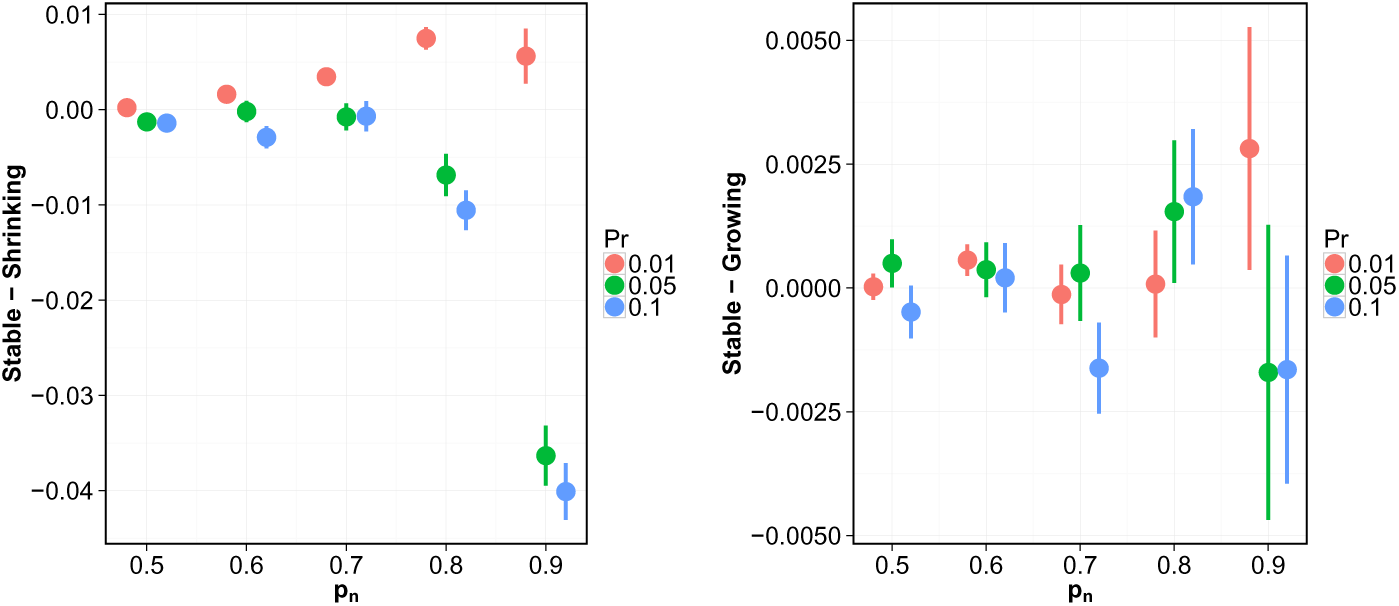
Density of growing and shrinking networks. The difference in network density of simulated networks from our model between stable and shrinking (left) or growing (right) networks. Points and lines represent the mean difference and standard error, respectively

**SI Figure 13:**
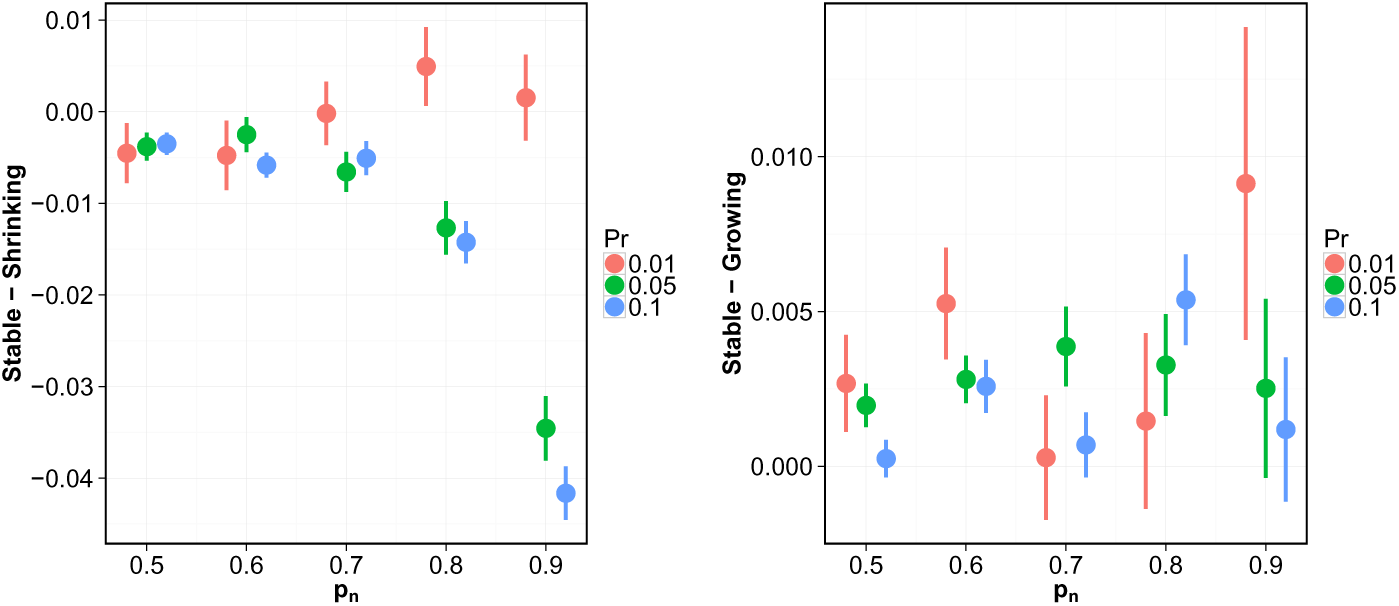
Clustering coefficients of growing and shrinking networks. The difference in global clustering coefficient of simulated networks from our model between stable and shrinking (left) or growing (right) networks. Points and lines represent the mean difference and standard error, respectively.

**SI Figure 14:**
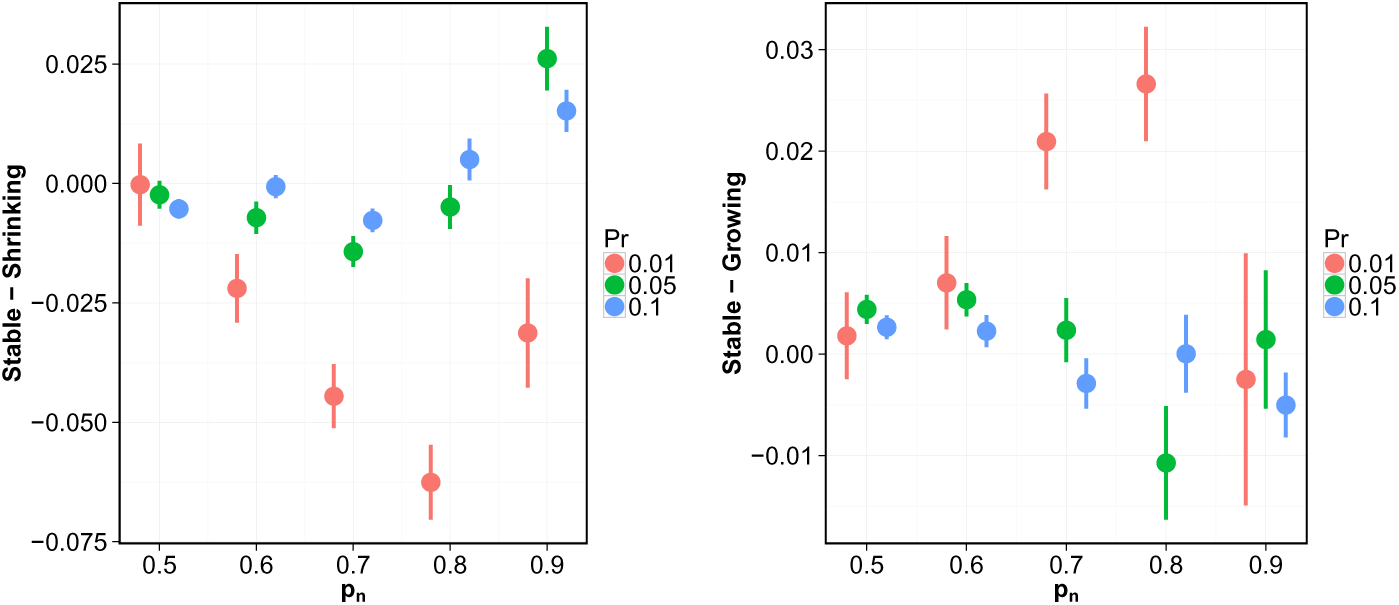
Modularity of growing and shrinking networks. The difference in network modularity of simulated networks from our model between stable and shrinking (left) or growing (right) networks. Points and lines represent the mean difference and standard error, respectively.

The effect of shrinking the network size was not consistent for all parameter sets. Nevertheless, shrinking networks tended to be denser in ties and less modular than networks of constant size for low *p_r_*. In a similar fashion, the effect of growing network size was not consistent for all parameter sets.

We conclude that the effect of changes in population size on network structure is unpredictable, and depends on the bonding probabilities. Future work should explore many interesting questions about the interaction of population size and social structure.

